# Degeneracy in the robust expression of spectral selectivity, subthreshold oscillations and intrinsic excitability of entorhinal stellate cells

**DOI:** 10.1101/197392

**Authors:** Divyansh Mittal, Rishikesh Narayanan

## Abstract

Biological heterogeneities are ubiquitous and play critical roles in the emergence of physiology at multiple scales. Although neurons in layer II (LII) of the medial entorhinal cortex (MEC) express heterogeneities in their channel properties, the impact of such heterogeneities on the robustness of cellular-scale physiology has not been assessed. Here, we performed a 55-parameter stochastic search spanning 9 voltage- or calcium-activated channels to assess the impact of channel heterogeneities on the concomitant emergence of 10 electrophysiological characteristics of LII stellate cells (SCs). We generated 50,000 models and found a heterogeneous subpopulation of 155 valid models to robustly match all electrophysiological signatures. We employed this heterogeneous population to demonstrate the emergence of cellular-scale degeneracy in LII SCs, whereby disparate parametric combinations expressing weak pairwise correlations resulted in similar models. We then assessed the impact of virtually knocking out each channel from all valid models and demonstrate that the mapping between channels and measurements was many-to-many, a critical requirement for the expression of degeneracy. Finally, we quantitatively predict that the spike-triggered average of LII SCs should be endowed with theta-frequency spectral selectivity and coincidence detection capabilities in the fast gamma-band. We postulate this fast gamma-band coincidence detection as an instance of cellular-scale efficient coding, whereby SC response characteristics match the dominant oscillatory signals in LII MEC. The heterogeneous population of valid SC models built here unveils the robust emergence of cellular-scale physiology despite significant channel heterogeneities, and forms an efficacious substrate for evaluating the impact of biological heterogeneities on entorhinal network function.

**KEY POINTS:**

- Stellate cells (SC) in layer II (LII) of the medial entorhinal cortex express cellular-scale degeneracy in the concomitant manifestation of several of their unique physiological signatures.
- Several disparate parametric combinations expressing weak pairwise correlations resulted in models with very similar physiological characteristics, including robust theta-frequency membrane potential oscillations spanning several levels of subthreshold depolarization.
- Electrophysiological measurements of LII SCs exhibited differential and variable dependencies on underlying channels, and the mapping between channels and measurements was many-to-many.
- Quantitative predictions point to theta-frequency spectral selectivity and fast gamma-range coincidence detection capabilities in class II/III spike-triggered average of LII SCs, with the postulate for this to be an instance of cellular-scale efficient coding.
- A heterogeneous cell population that accounts for both channel and intrinsic heterogeneities in LII SCs, which could be employed by network models of entorhinal function to probe the impact of several biological heterogeneities on spatial navigation circuits.

## INTRODUCTION

Networks in the nervous system are endowed with several forms of heterogeneities, which are known to play vital roles in the emergence of physiology and behavior. These ubiquitous forms of heterogeneities have been shown to either aid or hamper physiology in a manner that is reliant on several variables including the system under consideration, its specific function and the state of the system. Such disparate state-dependent impact of biological heterogeneities have necessitated system- and state-dependent quantitative analyses in assessing the precise role of these heterogeneities in specific neuronal structures and associated emergent functions (Wang & Buzsaki, 1996; Prinz *et al.*, 2004; Marder & Goaillard, 2006; Shamir & Sompolinsky, 2006; Chelaru & Dragoi, 2008; Goaillard *et al.*, 2009; Nusser, 2009; Grashow *et al.*, 2010; Padmanabhan & Urban, 2010; Ecker *et al.*, 2011; Marder, 2011; Marder & Taylor, 2011; Angelo *et al.*, 2012; Rathour & Narayanan, 2012; Tripathy *et al.*, 2013; Zhou *et al.*, 2013; Marder *et al.*, 2014; Rathour & Narayanan, 2014; Voliotis *et al.*, 2014; Anirudhan & Narayanan, 2015; Srikanth & Narayanan, 2015; Tikidji-Hamburyan *et al.*, 2015; Cadwell *et al.*, 2016; Fuzik *et al.*, 2016; Gjorgjieva *et al.*, 2016; Kohn *et al.*, 2016; Das *et al.*, 2017; Mukunda & Narayanan, 2017).

Neurons in layer II (LII) of the rodent medial entorhinal cortex (MEC) have been implicated in spatial navigation, especially since the cells in LII MEC are known to act as grid cells that elicit action potentials in a grid-like pattern as the animal traverses an arena (Hafting *et al.*, 2005; Moser *et al.*, 2008; Buzsaki & Moser, 2013; Moser *et al.*, 2014; Ray *et al.*, 2014; Tang *et al.*, 2014; Moser *et al.*, 2015). Ever since the discovery of grid cells, several theoretical and computational models have been proposed for their emergence, and have been tested from several different perspectives with varying degrees of success (O'Keefe & Burgess, 2005; Burak & Fiete, 2006; Burgess *et al.*, 2007; Jeewajee *et al.*, 2008; Welinder *et al.*, 2008; Burak & Fiete, 2009; Giocomo *et al.*, 2011b; Sreenivasan & Fiete, 2011; Navratilova *et al.*, 2012; Couey *et al.*, 2013; Domnisoru *et al.*, 2013; Schmidt-Hieber & Hausser, 2013; Yoon *et al.*, 2013; Bush & Burgess, 2014; Rowland *et al.*, 2016; Schmidt-Hieber *et al.*, 2017). Although these models and associated experiments have provided significant insights into entorhinal function, a lacuna common to these models relates to the systematic assessment of the impact of the different biological heterogeneities in the medial entorhinal cortex. Specifically, a systematic evaluation of the role of different forms of network heterogeneities, including those in channels, structural, intrinsic and synaptic properties and in afferent connectivity, with reference to entorhinal physiology has been lacking.

A first and essential step in addressing these and other related questions on the impact of biological heterogeneities on entorhinal network function is to assess the robustness of cellular physiology in the presence of well-established heterogeneities in channel properties. Specifically, measurements of channel properties, including kinetics, voltage-dependent gating and conductance values, from entorhinal neurons are known to exhibit significant variability across neurons (Bruehl & Wadman, 1999; Magistretti & Alonso, 1999; Dickson *et al.*, 2000; Fransen *et al.*, 2004; Castelli & Magistretti, 2006; Dudman & Nolan, 2009; Pastoll *et al.*, 2012; Schmidt-Hieber & Hausser, 2013). Despite this, entorhinal neurons exhibit signature electrophysiological characteristics that robustly fall into specific ranges for each physiologically relevant measurement. How do these neurons achieve such cellular-scale robustness in their physiology despite widely variable conductances and channel properties? Is there a requirement for individual channels or pairs of channels to be maintained at specific levels with specific properties for signature electrophysiological properties to emerge? Is there a one-to-one mapping between individual channels and the physiological properties that they regulate?

In this study, we build a conductance-based intrinsically heterogeneous population of LII stellate cell (SC) models of the medial entorhinal cortex that satisfied several of their unique electrophysiological signatures. We employed this heterogeneous population of LII SC models to demonstrate the expression of cellular-scale degeneracy (Edelman & Gally, 2001) in the concomitant emergence of these measurements. Specifically, we showed that LII SC with very similar electrophysiological characteristics emerged from disparate channel and parametric combinations expressing weak pairwise correlations. We employed these models to demonstrate the differential and variable dependencies of measurements on underlying channels, and showed that the mapping between channels and measurements was many-to-many. Finally, we employed this electrophysiologically validated model population to make quantitative testable predictions that the spike triggered average (STA) of LII SCs should be endowed with theta-frequency spectral selectivity and coincidence detection capabilities in the fast gamma-band. We postulate this fast gamma-band coincidence detection to be an instance of cellular-scale efficient coding (Narayanan & Johnston, 2012) that matches neuronal response properties to the dominant oscillatory band in the superficial layers of MEC (Colgin *et al.*, 2009; Colgin & Moser, 2010; Colgin, 2016; Trimper *et al.*, 2017). The heterogeneous population of models built here also forms an efficacious substrate for constructing network models of the entorhinal cortex, towards assessing the impact of cellular and channel properties and associated heterogeneities on emergent behavior such as grid cell formation.

## METHODS

We employed a single compartmental cylinder model of 70-µm diameter and 75-µm length (Fig. 1A). The choice of a single compartmental model was largely driven by the absence of direct and detailed electrophysiological characterization of dendritic intrinsic properties or of ion channels that express in LII SCs. As a consequence, morphologically precise models with specific channel expression profiles and matched intrinsic properties were infeasible. On the other hand, as the somatic channel properties and intrinsic physiological measurements of LII SCs are well characterized, we employed a single-compartmental model that wouldn't have to make explicit or implicit assumptions about dendritic physiology. Additionally, as a goal of this study was to develop an intrinsic heterogeneous model population of LII SCs towards their incorporation into network models, it was essential to ensure that the computational complexity of single neurons was minimal. A single compartmental conductance-based model population that was endowed with the different ion channels and systematically reflects intrinsic heterogeneities in LII SCs served as an efficacious means to achieve this goal as well.

**Figure 1:**
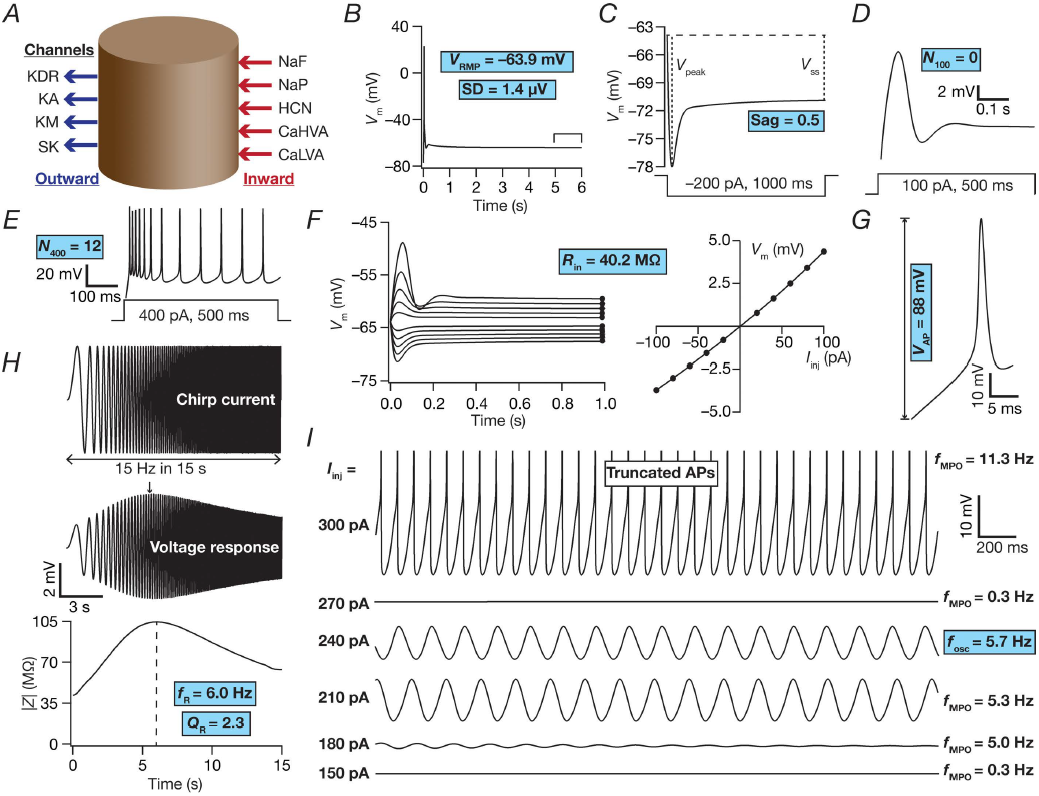
Base model and measurements. (*A*) Schematic representation of a single compartment model for MEC layer II stellate cell specifying inward and outward currents. (*B–I*) The 10 physiologically relevant measurements (highlighted in cyan) used to characterize stellate cells. (*B*) Resting membrane potential (*V*_RMP_) and its standard deviation (SD) were computed by taking the mean and standard deviation, respectively, of the membrane potential between 5–6 s duration (window specified in the figure) when no current was injected. All the other measurements were performed after the model settled at its *V*_RMP_ at 6 s. (*C*) Sag ratio (Sag) was measured as the ratio of the steady-state membrane potential deflection (*V*_SS_) to peak membrane potential deflection (*V*_peak_) in the voltage response of the model to a hyperpolarizing step current of 200 pA for a duration of 1000 ms. (*D–E*) Voltage response of the model to a step current of 100 pA (*D*) or 400 pA (*E*) for a stimulus duration of 500 ms was used to measure the number of action potentials (*N*_100_ or *N*_400_) elicited for the respective current injection. (*F*) Input resistance (*R*_in_) computation. *Left*, 1000 ms long step currents from –100 pA to 100 pA were injected into the cell in steps of 20 pA to record the steady state voltage response (black circles at the end of each trace). *Right*, Steady-state voltage response *vs.* injected current (*V–I*) plot obtained from the traces on the left panel. The slope of a linear fit to the *V–I* plot defined *R*_in_. (*G*) Amplitude of action potential (*V*_AP_) was measured as the difference between the peak voltage achieved during an action potential and *V*_RMP_. (*H*) Impedance based measurements. *Top*, Chirp current stimulus injected into the cell. *Middle*, Voltage response of the model to chirp stimulus injection. The arrow depicts the location of the maximal response. *Bottom*, Impedance amplitude profile showing the resonance frequency (*f*_R_) at which the model elicited peak response and resonance strength (*Q*_R_), the ratio of impedance amplitude at *f*_R_ to impedance amplitude at 0.5 Hz. (*I*) Membrane potential oscillations (MPOs). Shown are representative voltage traces (3 s duration) for different depolarizing current injections (*I*_inj_). The emergence of subthreshold oscillations in the theta range may be observed in traces at intermediate values of *I*_inj_, with the model switching to action potential firing at higher *I*_inj_. The frequency of subthreshold oscillations measured at a peri-threshold voltage was defined as *f*_osc_, while the frequency of membrane potential oscillations obtained with other *I*_inj_ was represented by *f*_MPO_.

### Passive and active neuronal properties

Passive properties were incorporated into the model as RC circuit that was defined through a specific membrane resistance, *R*_m_ and a specific membrane capacitance, *C*_m_. We introduced 9 different active channel conductances into the model (Fig. 1A): fast sodium (NaF), delayed rectifier potassium (KDR), hyperpolarization-activated cyclic-nucleotide gated (HCN) nonspecific cationic, persistent sodium (NaP), *A*-type potassium (KA), low-voltage activated calcium (LVA), high-voltage activated calcium (HVA), *M-*type potassium (KM) and small conductance calcium gated potassium (SK) channels. The channel kinetics and voltage dependencies for NaF, KDR and KA were from (Dudman & Nolan, 2009), for HCN from (Dickson *et al.*, 2000; Fransen *et al.*, 2004; Schmidt-Hieber & Hausser, 2013), for NaP from (Magistretti & Alonso, 1999; Dickson *et al.*, 2000; Fransen *et al.*, 2004), for HVA from (Bruehl & Wadman, 1999; Castelli & Magistretti, 2006), for LVA from (Bruehl & Wadman, 1999; Pastoll *et al.*, 2012), for SK from (Sah & Isaacson, 1995; Sah & Clements, 1999) and for KM from (Shah *et al.*, 2008).

All channel models were based on Hodgkin-Huxley formulation (Hodgkin & Huxley, 1952) except for the SK channel, which was modeled using a six-state Markovian kinetics model. Sodium, potassium and HCN channel currents were modeled using the Ohmic formulation and calcium channels followed the Goldman-Hodgkin-Katz (GHK) formulation (Goldman, 1943; Hodgkin & Katz, 1949) for current computation. The reversal potentials for Na^+^, K^+^ and HCN channel were 50, –90 and –20 mV, respectively. Calcium current through voltage gated calcium channels contributed to cytosolic calcium concentration ([*Ca*]_*c*_), and its decay was defined through simple first order kinetics (Destexhe *et al.*, 1993; Poirazi *et al.*, 2003; Carnevale & Hines, 2006; Narayanan & Johnston, 2010; Honnuraiah & Narayanan, 2013):

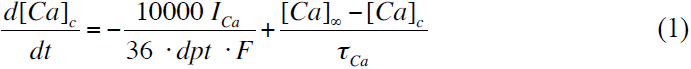

where *F* represented Faraday's constant, τ_ca_ defined the calcium decay time constant, *dpt* = 0.1 µm was the depth of the shell for cytosolic calcium dynamics and [*Ca*]_∞_ = 100 nM, was the steady state value of the [*Ca*]_*c*_.

Channel models were directly adopted in cases where they were explicitly based on direct measurements from LII SC channels. In cases where such explicit models were not available, model formulations were taken from other cell types and were explicitly adapted to match direct electrophysiological measurements. As channel models were either adopted from different studies or were adapted to match experimental observation, in what follows, we provide details of the models that we employed for gating and kinetics of each channel. The parameters that define these channel models are described in Table 1, along with the base values of the 55 passive and active parameters that govern these models. In channels that employed the Hodgkin-Huxley formulation, the model evolved by modifications to one or two gating particles, with each gating particle following first-order kinetics as follows:

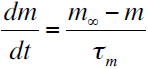

where *m*_∞_ and τ_*m*_ respectively defined the steady-state value and the time constant of the state variable that governed the gating particle. Channel gating and kinetics were appropriately adjusted for temperature dependence from corresponding experimental measurements.

**Table 1:** Base value and range of parameters used in generating the model population

| No | Parameter | Description | Base | Min | Max |
| --- | --- | --- | --- | --- | --- |
| **Fast sodium (NaF) channel** |  |  |  |  |  |
| 1 | $g^{NaF}$ (mS/cm<sup>2</sup>) | Maximal conductance of NaF | 4.2 | 2.1 | 8.5 |
| 2 | $V_m^{NaF}$ (mV) | Half-maximal voltage of activation of NaF | -26.1 | -31.1 | -21.1 |
| 3 | $k_m^{NaF}$ (mV) | Slope of activation of NaF | 9.38 | 7.51 | 11.26 |
| 4 | $F_m^{NaF}$ | Scaling factor for activation time constant of NaF | 1 | 0.8 | 1.2 |
| 5 | $V_h^{NaF}$ (mV) | Half-maximal voltage of inactivation of NaF | -23.8 | -28.8 | -18.8 |
| 6 | $k_h^{NaF}$ (mV) | Slope of inactivation of NaF | 6.1 | 4.9 | 7.3 |
| 7 | $F_h^{NaF}$ | Scaling factor for inactivation time constant of NaF | 1 | 0.8 | 1.2 |
| **Delayed rectifier potassium (KDR) channel** |  |  |  |  |  |
| 8 | $g^{KDR}$ (mS/cm<sup>2</sup>) | Maximal conductance of KDR | 3.2 | 1.5 | 6.4 |
| 9 | $V_n^{KDR}$ (mV) | Half-maximal voltage of activation of KDR | -17.6 | -22.6 | -12.6 |
| 10 | $k_n^{KDR}$ (mV) | Slope of activation of KDR | 19.6 | 15.7 | 23.6 |
| 11 | $F_n^{KDR}$ | Scaling factor for activation time constant of KDR | 1 | 0.8 | 1.2 |
| **Hyperpolarization-activated cyclic-nucleotide-gated (HCN) channel** |  |  |  |  |  |
| 12 | $g^{HCN}$ (μS/cm<sup>2</sup>) | Maximal conductance of slow HCN | 33.3 | 16 | 67 |
| 13 | $HCN_{ms}^{mf}$ | Ratio of fast to slow HCN maximal conductance | 1.85 | 1.5 | 2.2 |
| 14 | $V_{mf}^{HCN}$ (mV) | Half-maximal voltage of activation of fast HCN | 74.2 | 69.2 | 79.2 |
| 15 | $V_{ms}^{HCN}$ (mV) | Half-maximal voltage of activation of slow HCN | 2.83 | -2.17 | 7.83 |
| 16 | $k_{mf}^{HCN}$ (mV) | Slope of activation of fast HCN | 9.78 | 7.8 | 11.7 |
| 17 | $k_{ms}^{HCN}$ (mV) | Slope of activation of slow HCN | 15.9 | 12.7 | 19.1 |
| 18 | $F_{mf}^{HCN}$ | Scaling factor for activation time constant of fast HCN | 1 | 0.8 | 1.2 |
| 19 | $F_{ms}^{HCN}$ | Scaling factor for activation time constant of slow HCN | 1 | 0.8 | 1.2 |
| **Persistent sodium (NaP) channel** |  |  |  |  |  |
| 20 | $g^{NaP}$ (μS/cm<sup>2</sup>) | Maximal conductance of NaP | 34 | 17 | 68 |
| 21 | $V_m^{NaP}$ (mV) | Half-maximal voltage of activation of NaP | 48.7 | 43.7 | 53.7 |
| 22 | $k_m^{NaP}$ (mV) | Slope of activation of NaP | 4.4 | 3.52 | 5.28 |
| 23 | $F_m^{NaP}$ | Scaling factor for activation time constant of NaP | 1 | 0.8 | 1.2 |
| 24 | $V_h^{NaP}$ (mV) | Half-maximal voltage of inactivation of NaP | 48.8 | 43.8 | 53.8 |
| 25 | $k_h^{NaP}$ (mV) | Slope of inactivation of NaP | 9.9 | 7.9 | 11.9 |
| 26 | $F_h^{NaP}$ | Scaling factor for inactivation time constant of NaP | 1 | 0.8 | 1.2 |

| **A-type potassium (KA) channel** |  |  |  |  |  |
| --- | --- | --- | --- | --- | --- |
| 27 | $g^{KA}$ ($\mu\text{S}/\text{cm}^2$) | Maximal conductance of KA | 25 | 12.5 | 50 |
| 28 | $V_m^{KA}$ (mV) | Half-maximal voltage of activation of KA | -18.3 | -23.3 | -13.3 |
| 29 | $k_m^{KA}$ (mV) | Slope of activation of KA | 15 | 12 | 18 |
| 30 | $F_m^{KA}$ | Scaling factor for activation time constant of KA | 1 | 0.8 | 1.2 |
| 31 | $V_h^{KA}$ (mV) | Half-maximal voltage of inactivation of KA | -58 | -63 | -53 |
| 32 | $k_h^{KA}$ (mV) | Slope of inactivation of KA | 8.2 | 6.6 | 9.8 |
| 33 | $F_h^{KA}$ | Scaling factor for inactivation time constant of KA | 1 | 0.8 | 1.2 |
| **High voltage activated (HVA) calcium channel** |  |  |  |  |  |
| 34 | $g^{HVA}$ ($\text{mS}/\text{cm}^2$) | Maximal conductance of HVA | 0.18 | 0.09 | 0.36 |
| 35 | $V_m^{HVA}$ (mV) | Half-maximal voltage of activation of HVA | 11.1 | 6.1 | 16.1 |
| 36 | $k_m^{HVA}$ (mV) | Slope of activation of HVA | 8.4 | 6.7 | 10.0 |
| 37 | $F_m^{HVA}$ | Scaling factor for activation time constant of HVA | 1 | 0.8 | 1.2 |
| 38 | $V_h^{HVA}$ (mV) | Half-maximal voltage of inactivation of HVA | 37 | 32 | 42 |
| 39 | $k_h^{HVA}$ (mV) | Slope of inactivation of HVA | 9 | 7.2 | 10.8 |
| 40 | $F_h^{HVA}$ | Scaling factor for inactivation time constant of HVA | 1 | 0.8 | 1.2 |
| **Low voltage activated (LVA) calcium channel** |  |  |  |  |  |
| 41 | $g^{LVA}$ ($\mu\text{S}/\text{cm}^2$) | Maximal conductance of LVA | 90 | 41.9 | 167.6 |
| 42 | $V_m^{LVA}$ (mV) | Half-maximal voltage of activation of LVA | -52.4 | -57.4 | -47.4 |
| 43 | $k_m^{LVA}$ (mV) | Slope of activation of LVA | 8.2 | 6.5 | 9.8 |
| 44 | $F_m^{LVA}$ | Scaling factor for activation time constant LVA | 1 | 0.8 | 1.2 |
| 45 | $V_h^{LVA}$ (mV) | Half-maximal voltage of inactivation of LVA | -88.2 | -93.2 | -83.2 |
| 46 | $k_h^{LVA}$ (mV) | Slope of inactivation of LVA | 6.67 | 5.34 | 8.01 |
| 47 | $F_h^{LVA}$ | Scaling factor for inactivation time constant of LVA | 1 | 0.8 | 1.2 |
| **M-type potassium (KM) channel** |  |  |  |  |  |
| 48 | $g^{KM}$ ($\text{mS}/\text{cm}^2$) | Maximal conductance of KM | 0.12 | 0.06 | 0.25 |
| 49 | $V_m^{KM}$ (mV) | Half-maximal voltage of activation of KM | -40 | -45 | -35 |
| 50 | $k_m^{KM}$ (mV) | Slope of activation of KM | -10 | -8 | -12 |
| 51 | $F_m^{KM}$ | Scaling factor for activation time constant of KM | 1 | 0.8 | 1.2 |
| **Small conductance calcium-activated potassium (SK) channel** |  |  |  |  |  |
| 52 | $g^{SK}$ ($\mu\text{S}/\text{cm}^2$) | Maximal conductance of SK | 52 | 26 | 104 |

| Passive properties and cytosolic calcium handling |  |  |  |  |  |
| --- | --- | --- | --- | --- | --- |
| 53 | $R_m$ (kΩ cm<sup>2</sup>) | Specific membrane resistance | 40 | 20 | 80 |
| 54 | $\tau_{Ca}$ (ms) | Time constant of cytosolic calcium decay | 78 | 39 | 156 |
| 55 | $C_m$ (μF/cm<sup>2</sup>) | Specific membrane capacitance | 1 | 0.75 | 1.25 |

#### The fast sodium channel

The NaF model was adopted from (Dudman & Nolan, 2009), and the current through this sodium channel was:

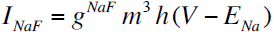

The activation gating particle was defined by:

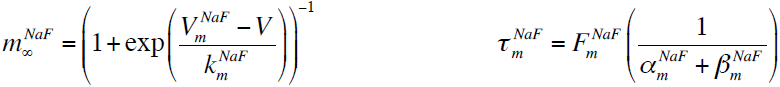

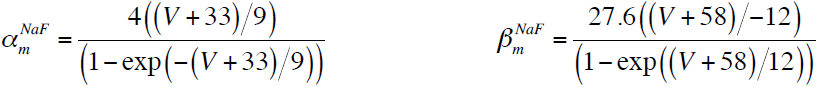

The inactivation gating particle was defined by:

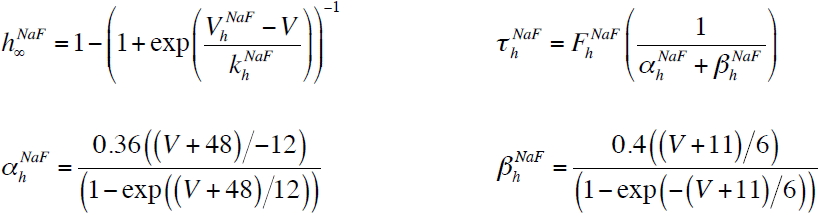

#### The delayed rectifier potassium channel

The KDR model was adopted from (Dudman & Nolan, 2009), and the current through this potassium channel was:

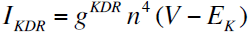

The activation gating particle was governed by the following equations:

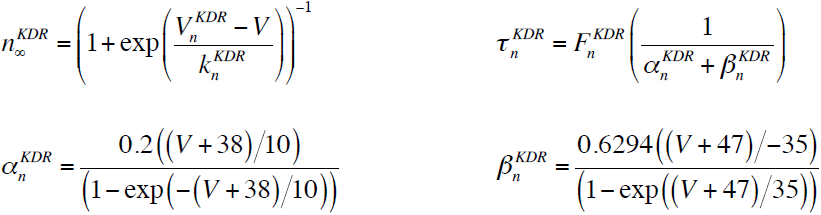

#### The hyperpolarization-activated cyclic-nucleotide-gated channel

The HCN channel model was adopted from (Dickson *et al.*, 2000; Fransen *et al.*, 2004; Schmidt-Hieber & Hausser, 2013) and the current through this nonspecific cationic channel was:

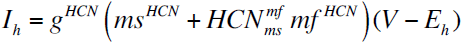

where *ms*^*HCN*^ and *mf*^*HCN*^ respectively defined the gating variables for the slow and fast components of the current through HCN channels, and 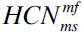 defined the ratio of the fast to slow HCN conductance values. The activation gating particles for the slow and fast HCN components were governed by the following equations:

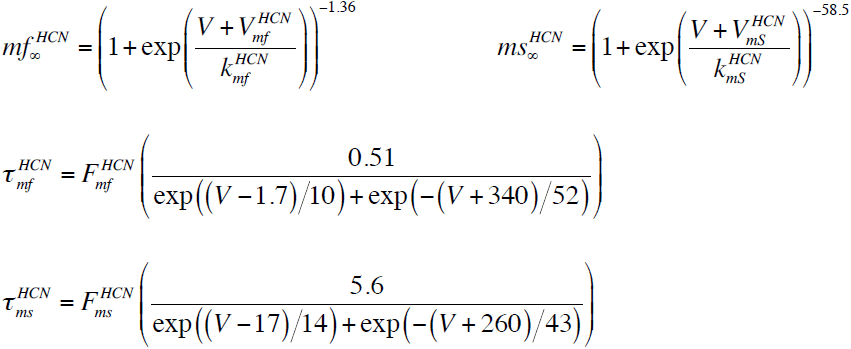

#### The persistent sodium channel

The NaP model was adopted from (Magistretti & Alonso, 1999; Dickson *et al.*, 2000; Fransen *et al.*, 2004), and the current through this sodium channel was:

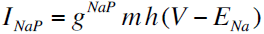

The activation gating particle was defined by:

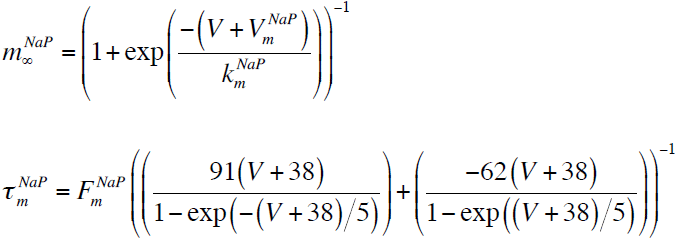

The inactivation gating particle was defined by:

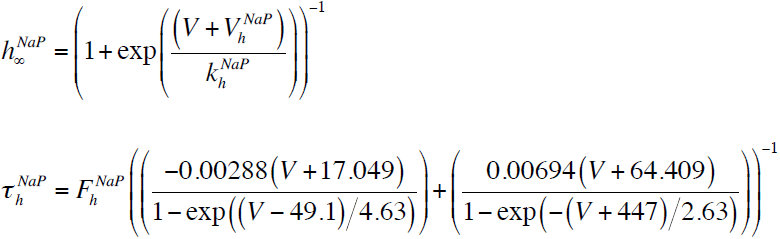

#### The transient A-type potassium channel

The KA model was adopted from (Dudman & Nolan, 2009), and the current through this potassium channel was

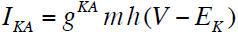

The activation gating particle was governed by the following equations:

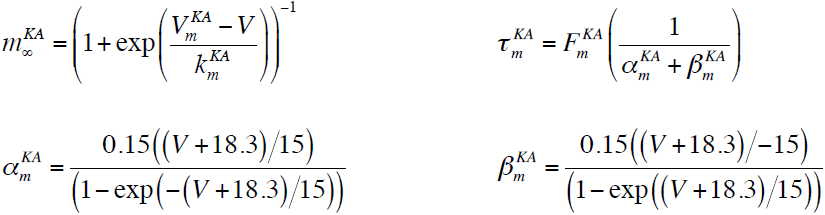

The inactivation gating particle was governed by the following equations:

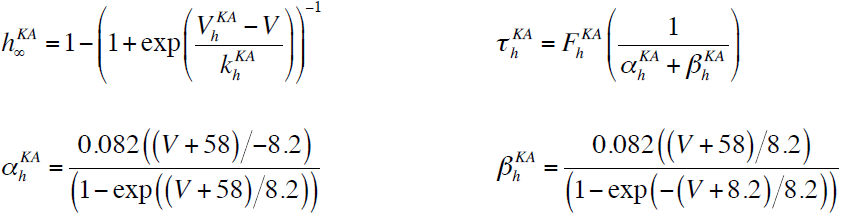

#### The high voltage-activated calcium channel

The HVA calcium channel model was fitted with corresponding electrophysiological data (Bruehl & Wadman, 1999; Castelli & Magistretti, 2006). The current through this channel followed GHK conventions, with the default extracellular and cytosolic calcium concentrations set at 2 mM and 100 nM, respectively. The conductance evolution of this channel was modeled as follows:

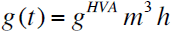

The activation and inactivation gating particles were governed by the following equations:

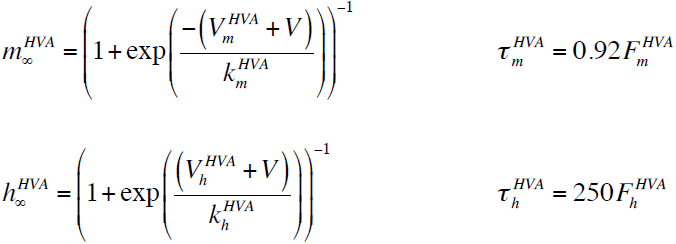

#### The low voltage-activated calcium channel

The LVA calcium channel model was fitted with corresponding electrophysiological data from (Bruehl & Wadman, 1999; Pastoll *et al.*, 2012). The current through this channel followed GHK conventions, with the default extracellular and cytosolic calcium concentrations set at 2 mM and 100 nM, respectively. The conductance evolution of this channel was modeled as follows:

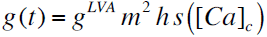

where *m* and *h* respectively represented the voltage-dependent activation and inactivation gating particles, and *s* ([*Ca*]_*c*_)governed calcium-dependent inactivation with [*Ca*]_*c*_ specified in mM. Their evolution was dictated by the following equations:

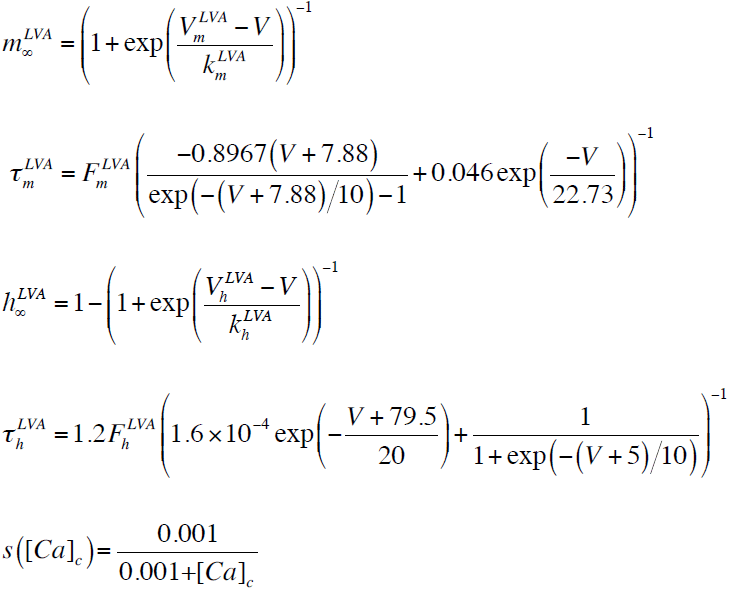

#### The M-type potassium channel

The KM model was adopted from (Shah *et al.*, 2008), and the current through this potassium channel was:

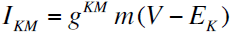

The activation gating particle was governed by the following equations:

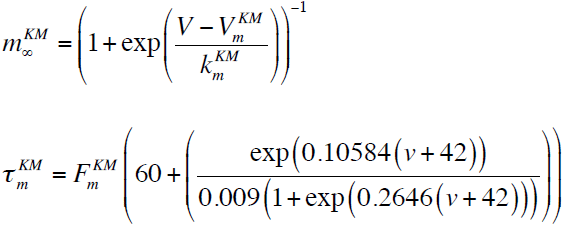

### Intrinsic measurements

We measured the resting membrane potential of the model neuron by allowing the model to settle at a steady-state potential when no current was injected for a period of 5 s. This was essential because there were several slow subthreshold conductances (especially the calcium-activated potassium channel) that contributed to resting membrane potential in our model. We set the passive membrane potential, in the absence of any subthreshold conductance, to be at –77 mV (Boehlen *et al.*, 2013) and allowed the interactions among the several subthreshold conductances to set the steady-state resting membrane potential (*V*_RMP_). After this initial 5 s period of the simulation, *V*_RMP_ was computed as the mean of the membrane potential over a one-second interval (5^th^ to 6^th^ second; Fig. 1B). We also calculated the standard deviation (SD) of membrane potential over the same 1 s period to ensure that the RMP was measured after attainment of steady state. All intrinsic measurements reported below were always performed *after* this 6 s period (5 s for the transients to settle to steady-state and 1 s for RMP measurement).

To estimate sag ratio in the model, we injected a hyperpolarizing step current of 200 pA amplitude for 1s and recorded the voltage response. Sag ratio (Sag) was computed as the membrane potential deflection achieved at steady state (*V*_SS_) during the current injection period divided by the peak deflection of the membrane potential (*V*_peak_) within the period of current injection (Fig. 1C). In assessing supra-threshold excitability of the model neuron, we measured the number of action potentials (AP) elicited by the neuron in response to different depolarizing step current injections spanning 500 ms. We defined the number of APs fired for 100 and 400 pA current injections as *N*_100_ (Fig. 1D) and *N*_400_ (Fig. 1E), respectively.

Input resistance (*R*_in_) was calculated from the steady-state voltage response (after 1 s of current injection) of the model neuron to subthreshold current pulses of amplitudes spanning –100 pA to 100 pA in steps of 20 pA. The steady state voltage response was plotted against the corresponding amplitude of injected current, and the slope of a linear fit to this plot was assigned as the input resistance of the model (Fig. 1F). Spike amplitude (*V*_AP_) was computed from the first AP elicited during a 400-pA step current injection, and was defined as the difference between *V*_RMP_ and the peak membrane potential achieved during the AP (Fig. 1G).

As entorhinal stellates reside within an oscillatory network, it was essential that excitability measures be computed in a frequency-dependent manner. To do this, we computed well-established impedance-based measurements from its amplitude and phase profiles (Hutcheon *et al.*, 1996a, b; Hutcheon & Yarom, 2000; Haas & White, 2002; Erchova *et al.*, 2004; Giocomo *et al.*, 2007; Narayanan & Johnston, 2007, 2008). These profiles were computed by measuring the voltage response of the model to a chirp stimulus, a sinusoidal current stimulus with constant amplitude (40 pA peak to peak amplitude) with frequency linearly spanning from 0–15 Hz in 15 s (Fig. 1H). Frequency-dependent impedance, *Z* (*f*), was computed as the ratio between the Fourier transform of this voltage response and the Fourier transform of the chirp stimulus. The magnitude of the complex quantity defined the impedance amplitude profile (Fig. 1H):

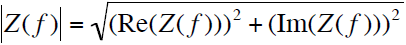

where Re(*Z*(*f*)) and Im(*Z*(*f*)) were the real and imaginary parts of the impedance *Z*(*f*), respectively. The frequency at which |*Z*(*f*)| reached its maximum value was considered as the resonance frequency, *f*_R_, and resonance strength (*Q*_R_) was defined as the ratio of |*Z*(*f*_R_)| to |*Z*(0.5)| (Fig. 1H). The impedance phase profile *Φ*(*f*) was computed as:

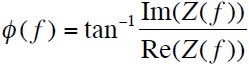

The total inductive area, Φ_L_, defined as the area under the inductive part of *Φ*(*f*), was calculated based on the impedance phase profile (Narayanan & Johnston, 2008):

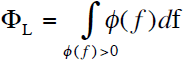

Sub- and peri-threshold membrane potential oscillations (MPO) were assessed in voltage responses of the model to depolarizing pulse current injections spanning 100–300 pA in steps of 10 pA, each lasting for 5 s (Fig. 1I). The last 3 s period of this 5-s period was transformed to frequency domain through the Fourier transform, and the frequency at which this spectral signal had maximum magnitude was defined as the MPO frequency (*f*_MPO_). We defined the peri-threshold oscillation frequency (*f*_osc_) as the *f*_MPO_ of the subthreshold voltage response proximal to the spiking threshold of the model.

### Multi-parametric Multi-objective Stochastic Search Algorithm

To generate an intrinsically heterogeneous population of LII SCs and to assess if the concomitant functional expression of all 10 intrinsic measurements manifested degeneracy in terms of the specific ion channel combinations that can elicit them, we employed a multi-parametric multi-objective stochastic search (MPMOSS) algorithm (Foster *et al.*, 1993; Goldman *et al.*, 2001; Prinz *et al.*, 2003; Marder & Taylor, 2011; Rathour & Narayanan, 2012, 2014; Anirudhan & Narayanan, 2015; Srikanth & Narayanan, 2015; Mukunda & Narayanan, 2017). This stochastic search was performed over 55 parameters (Table 1) and jointly validated against 10 sub- and supra-threshold measurements (Fig. 1; *V*_RMP_, SD, Sag ratio, *R*_in_, *f*_R_, *Q*_R_, *f*_osc_, *N*_100_, *N*_400_, *V*_AP_) towards matching electrophysiological recordings from LII SCs (Table 2). In executing the MPMOSS algorithm, we constructed a model neuron from specific values for each of the 55 parameters, each of which was independently and randomly picked from a uniform distribution whose bounds reflected the electrophysiological variability in that parameter (Table 1). For each such randomly chosen model, which ensured that we are not biasing our parametric ranges with any constraints, all 10 intrinsic measurements were computed and were compared against their respective electrophysiological bounds (Table 2). A model that satisfied *all* the 10 criteria for validation was declared valid. We repeated this procedure for 50,000 randomized picks of the 55 parameters, and validated these models against the 10 measurements. Intrinsic heterogeneity and degeneracy were then assessed in the resulting population of valid LII SC models and their parametric combinations. This assessment was performed through the analysis of parametric distributions and pairwise correlations among valid model parameters.

**Table 2:** Physiologically relevant range of LII stellate cell measurements

| No | Intrinsic measurement | Valid range |
| --- | --- | --- |
| 1 | Resting membrane potential, $V_{RMP}$ (mV) | −65 to −60 |
| 2 | Standard deviation of membrane potential (Resting), SD (mV) | <0.01 |
| 3 | Sag ratio | 0.35 to 0.65 |
| 4 | Input resistance, $R_{in}$ (MΩ) | 35 to 65 |
| 5 | Resonance strength, $Q_R$ | < 3.5 |
| 6 | Resonance frequency, $f_R$ (Hz) | 3 to 12 |
| 7 | Peri-threshold MPO frequency, $f_{osc}$ (Hz) | 3 to 12 |
| 8 | Number of APs for a 100 pA step current for 500 ms, $N_{100}$ | 0 |
| 9 | Number of APs for a 400 pA step current for 500 ms, $N_{400}$ | 7 to 16 |
| 10 | AP amplitude, $V_{AP}$ (mV) | >75 |

To assess the impact of individual channels on each of the 10 intrinsic measurements within the valid model population, we employed the virtual knockout model (VKM) approach (Rathour & Narayanan, 2014; Anirudhan & Narayanan, 2015; Mukunda & Narayanan, 2017). In doing this, we first set the conductance value of each of the 9 active ion channels independently to zero for each of the valid models. We then computed all the 10 intrinsic measurements for each model, and assessed the sensitivity of each measurement to the different channels from the statistics of post-knockout change in the measurements across all valid models. When some of the channels were knocked out, certain valid models elicited spontaneous spiking or showed depolarization-induced block (when depolarizing current were injected). These VKMs were not included into the analysis for assessing the sensitivities, because this precluded computation of all 10 measurements from such models.

### Spike triggered average and associated measurements

For estimation of STA, a zero-mean Gaussian white noise (GWN) with a standard deviation σ_noise_ was injected into the neuron for 1000 s. σ_noise_ was adjusted such that overall action potential firing rate was ˜1 Hz in the model under consideration. This ensured that the spikes were isolated and aperiodic, thereby establishing statistical independence of the current samples used in arriving at the STA (Aguera y Arcas & Fairhall, 2003; Das & Narayanan, 2014, 2015, 2017). The STA was computed from the injected current for a period of 300 ms preceding the spike and averaged over all spikes across the time period of simulation, translating to around 1000 spikes for each STA computation. STA kernels were smoothed using a median filter spanning a 1 ms window for representation purposes and for computing quantitative measurements that were derived from the STA.

Quantitative metrics for spectral selectivity in the STA, for coincidence detection windows and intrinsic excitability were derived from the STA and its Fourier transform (Das & Narayanan, 2014, 2015, 2017). Specifically, the frequency at which the magnitude of the Fourier transform of the STA peaked was defined as the STA characteristic frequency (*f*_STA_). STA selectivity strength (*Q*_STA_) was defined as the ratio of |STA(*f*_STA_)| to |STA(0.5)|. The peak positive current in the STA kernel was defined as 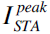, which constitutes a measure of excitability. To quantify the window for integration/coincidence detection, we defined the spike-proximal positive lobe (SPPL) as the temporal domain that was adjacent to the spike where the STA was positive (Das & Narayanan, 2015, 2017). The total coincidence detection window, CDW (*T*_TCDW_) was computed as the temporal distance from the spike location (*t*=0 ms) to the first zero crossing in the STA. *T*_TCDW_ constitutes the entire temporal expanse over which the inputs were positively weighted and hence covered the entire temporal spread of SPPL. To account for the specific shape of the STA in defining the coincidence detection window, we defined an effective CDW (*T*_ECDW_), which was a STA-weighted measure of SPPL (Das & Narayanan, 2015, 2017):

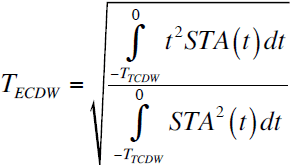

### Computational details

All simulations were performed using the NEURON programming environment (Carnevale & Hines, 2006) at 34° C, with a simulation step size of 25 µs. All data analyses and plotting were executed using custom-written software within the IGOR pro environment (Wavemetrics Inc.). All statistical analyses were performed using the R statistical package (R Core Team, 2013).

## RESULTS

The principal goal of this study was to assess the impact of molecular-scale heterogeneities in channel properties on cellular-scale physiological signatures of LII stellate cells of the medial entorhinal cortex. We approached this by building an unbiased stochastic search-based conductance-based population of LII SCs that satisfied several of their unique electrophysiological signatures. Apart from providing an efficacious substrate for understanding the roles of channel parameters, intrinsic measurements and associated heterogeneities in entorhinal function, our goal in building these models was three fold. First, a heterogeneous population of LII SC models that satisfied several electrophysiological constraints would provide us the means to assess if there was significant degeneracy in the emergence of these measurements, or if there was a requirement on unique mappings between channel properties and physiological measurements. Second, such a heterogeneous population would allow us to establish the specific roles of different channels in mediating or regulating different physiological properties, and assess variability in such regulatory roles. Third, and importantly, as these models match with LII SC models in several ways, they provide an ideal foundation for making quantitative predictions about stellate cell physiology, which can be electrophysiologically tested.

Towards this, we first hand-tuned a conductance-based, biophysically and physiologically relevant base model that was characterized as a single compartmental cylindrical model with different passive and active properties (see Methods). The model was endowed with 9 different active ion channels (besides leak channels), and matched with 10 distinct electrophysiological measurements obtained from LII SCs (Fig. 1, Table 2). These electrophysiological measurements included the significant sag observed in response to pulse current injections (Fig. 1C), theta-frequency membrane potential resonance that exhibited strong spectral selectivity (Fig. 1H), and importantly the robust subthreshold membrane potential oscillations in the theta-frequency range at different depolarized potentials (Fig. 1I). The 55 active and passive parameters that governed LII SC models, and their respective values in the base model are listed in Table 1.

### Diverse depolarization-dependent evolution of membrane potential oscillations in stochastically generated stellate cell models

We employed the base model parameters (Table 1) as the substrate for a multi-parametric stochastic search algorithm for models that would meet multiple objectives in terms of matching with the electrophysiological properties of LII SCs. The range of individual parameters over which this multi-parametric (55 parameters) multi-objective (10 measurements) stochastic search (MPMOSS) algorithm was executed is provided in Table 1. We generated a test population of 50,000 model cells by sampling these model parameters, and subjected these model cells to validation based on the physiologically observed range of sub- and supra-threshold measurements (Table 2). First, we found a subpopulation of these models where all measurements, except for the ability to express theta-frequency membrane potential oscillations, were within the specified bounds. We depolarized this subpopulation of models and asked if these models expressed robust subthreshold oscillations in their membrane potentials.

We found that the depolarization-dependent evolution of sub- and supra-threshold (regular spiking behavior) oscillations exhibited significant diversity across different models within this subpopulation (Fig. 2). In most models within this subpopulation, consistent with corresponding experimental observations (Alonso & Llinas, 1989), we observed the emergence of robust theta range subthreshold MPOs with membrane potential approaching near spiking threshold, with further depolarization resulting in spiking activity (Fig. 2A) or spike doublets (Fig. 2B) or bursts (Fig. 2C) riding over subthreshold MPOs. Among these, there were some models that exhibited theta skipping, where spikes or bursts were regular, but were not observed on every cycle of the subthreshold MPO (Fig. 2C). In other models, incrementally higher depolarization resulted in the cell switching from rest to subthreshold MPOs to a state that resembled depolarization-induced block (Fig. 2D–E). Whereas in certain models, further depolarization would result in regular spiking (Fig. 2D), in other models the depolarization-induced block persisted with the model never entering spiking behavior despite further depolarization (Fig. 2E). In rare cases where the model displayed spiking behavior without transitioning through subthreshold oscillations (Fig. 2F), the model was not included as a valid model because such models did not meet the electrophysiological constraint on theta-frequency peri-threshold oscillations. Finally, very few models manifested robust subthreshold oscillations, but switched back-and-forth between subthreshold oscillations and regular spiking with increasing current injections (Fig. 2G).

**Figure 2:**
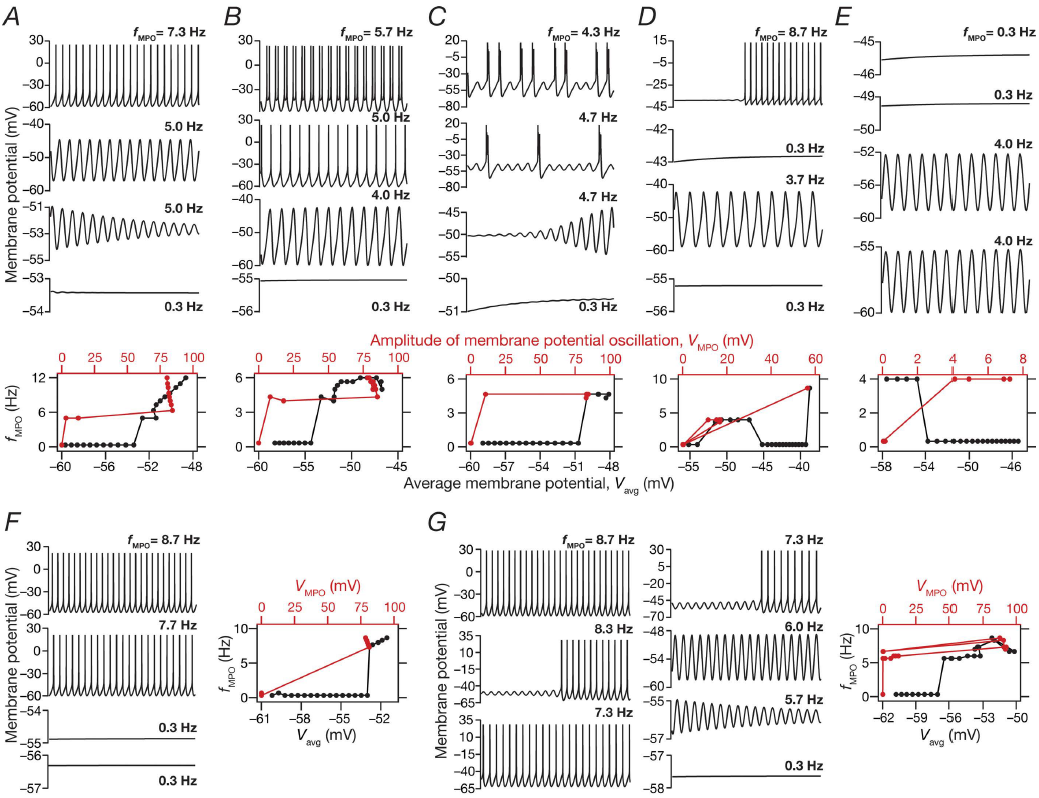
A multi-parametric stochastic search algorithm yielded stellate cell models with distinct types of robust sub- and supra-threshold oscillations spanning different voltage levels. (*A–G*) *Top*, Voltage traces of model cells showing sub and supra threshold MPOs, when injected with different levels of depolarizing currents. *Bottom (A–E) and Right (F–G)*, frequency of MPOs plotted as a function of average membrane potential, *V*_avg_ (black) and MPO amplitude, *V*_MPO_ (red). Plots in (*A–G*) constitute data from different model cells, and depict representative features from distinct subpopulations of models. (*A*) Robust subthreshold MPOs emerge before the neuron switches to regular spiking activity that manifests when the subthreshold MPOs cross threshold. (*B*) Robust subthreshold MPOs emerge before the neuron abruptly switches to firing spike doublets when the subthreshold MPOs cross threshold. (*C*) Neuron switches to robust MPOs at perithreshold voltages, with intermittent burst spiking activity. The frequency of burst occurrence increases with increasing current injections. Such models are reminiscent of neurons exhibiting theta skipping, where spikes occur at regular intervals but not on every theta cycle. (*D*) Model exhibits robust theta range subthreshold oscillations, but does not directly switch to spiking behavior from MPOs with increased current injection. A range of intermittent current injections results in responses that depict depolarization-induced block bereft of any MPOs. These models eventually switch to regular firing at higher current injections. (*E*) Same as (*D*) but these models do not switch to firing action potentials after exhibiting theta range MPOs even at higher depolarization or current injections. (*F*) These models abruptly switch from firing no action potentials to regular spiking, without any intermediate phase of exhibiting subthreshold oscillations. *(G)* Model manifests robust subthreshold oscillations, but switches between subthreshold oscillations and regular spiking with increasing current injections.

### The stochastic search strategy yielded an intrinsically heterogeneous population of models that matched several electrophysiological signatures of stellate cells

Out of 50,000 models that were generated as part of the stochastic search strategy, only 155 models (*N*_valid_ = 155) were valid when we constrained them against all the 10 electrophysiological measurements (Table 2), including their ability to manifest robust peri-threshold theta frequency oscillations. We plotted all the 10 electrophysiological measurements in this model population to assess if they were clustered around their respective base model values (Fig. 1) or if they were distributed to span the range of valid model measurements (Table. 2). Whereas a clustered set of measurements would have implied a near-homogeneous population of models, a distributed pattern that spanned the range of respective electrophysiological counterparts would provide us with a heterogeneous model population that reflects experimental variability in the respective measurements. We found all measurements to spread over a large span, with most of them covering the entire min-max range of their respective bounds (Fig. 3A; note that *N*_100_ has not been plotted because it was required to be zero for model validity). We observed the emergence of sub- and supra-threshold membrane potential oscillations in these models when the average membrane voltage was between –59 mV to –45 mV (Fig. 3B). To distinguish between sub- and supra-threshold oscillations, we plotted the frequency of these membrane potential oscillations against their peak-to-peak amplitude (Fig. 3C). Two clearly separable clusters were observed, with all subthreshold oscillations clustered at the low-amplitude range (< 25 mV), while the action potentials forming a cluster with amplitudes greater than ~60 mV. Importantly, these observations also demonstrate that the characteristics of membrane potential oscillations exhibited significant heterogeneity, thus matching the electrophysiological variability observed in LII SCs. Together, the biophysically and physiologically constrained MPMOSS algorithm yielded a population of LII SC models that manifested considerable intrinsic heterogeneities (Fig. 3) matching the ranges observed in corresponding electrophysiological measurements (Table 2).

**Figure 3:**
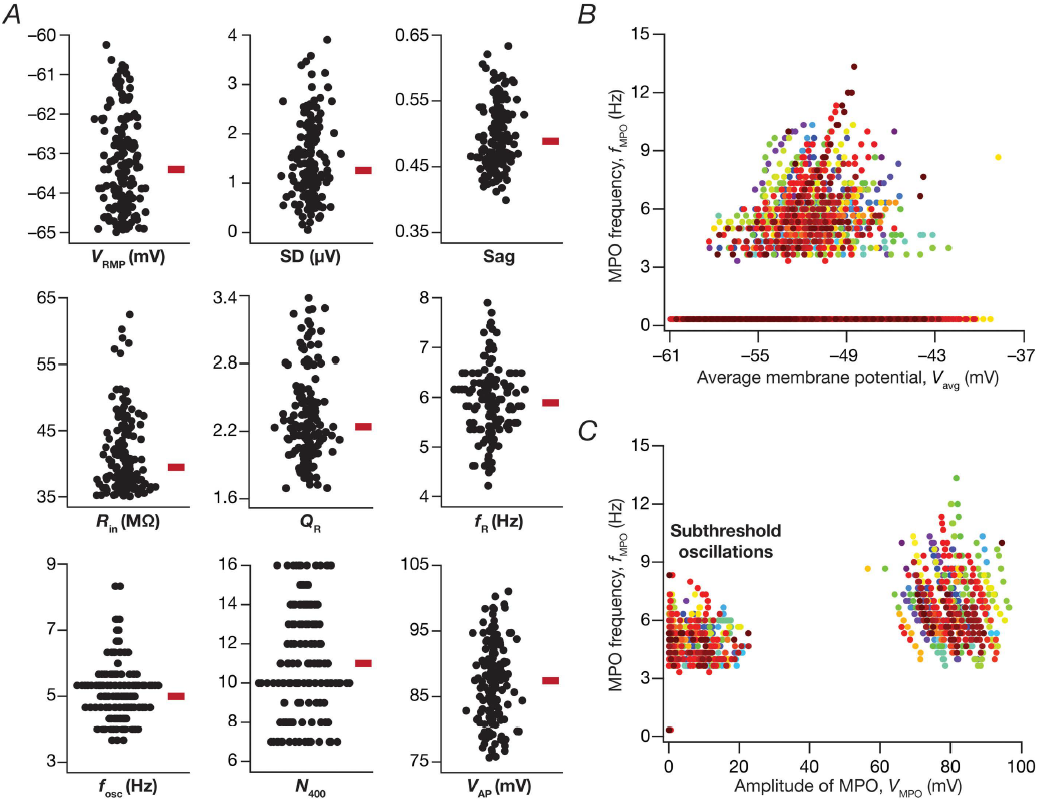
Distribution for physiologically relevant measurements in valid MEC layer II stellate cell models obtained after a multi-parametric multi-objective stochastic search. (*A*) Bee swarm plots depicting the distribution of 9 measurements in the 155 valid models. (*B*) Frequency of MPOs for the 155 valid models plotted as a function of average membrane potential of the oscillation, *V*_avg_. (*C*) Frequency of MPOs for the 155 valid models plotted as a function of MPO amplitude, *V*_MPO_. The two distinct clusters here demarcate sub- and supra-threshold oscillations, with supra-threshold oscillations corresponding to regular action potential firing. For *B–C*, 21 data points represent each valid model, with each data point obtained with different depolarizing current injections (*e.g.,* Fig. 2). Each model is depicted in a unique color.

### The valid model population manifested cellular-scale degeneracy

What were the specific constraints on the 55 different parameters in yielding the valid model population that *concomitantly* matched several electrophysiological signatures of stellate cells? Were these parameters clustered around specific values thereby placing significant constraints on the different channels, their properties and their expression profiles? Would individual channels have to be maintained at specific expression levels for models to match all 10 electrophysiological signatures? To address these questions, we first randomly picked 5 of the 155 valid models that exhibited similar measurements (Fig. 4A–H), and asked if the set of parameters governing these models were also similar. We normalized each of the 55 parameters from these 5 models with reference to their respective min-max values, and found none of these models to follow any specific trend in their parametric values with most parameters spreading across the entire range that they were allowed to span (Fig. 4I). These observations provided the first line of evidence for the expression of degeneracy in stellate cell models, whereby models with very similar functional characteristics emerged from disparate parametric combinations. To confirm this across all valid models, we plotted the histograms of each parameter for all the 155 valid models, and found their spread to span the entire testing range for all parameters (Fig. 5A, bottom row).

**Figure 4:**
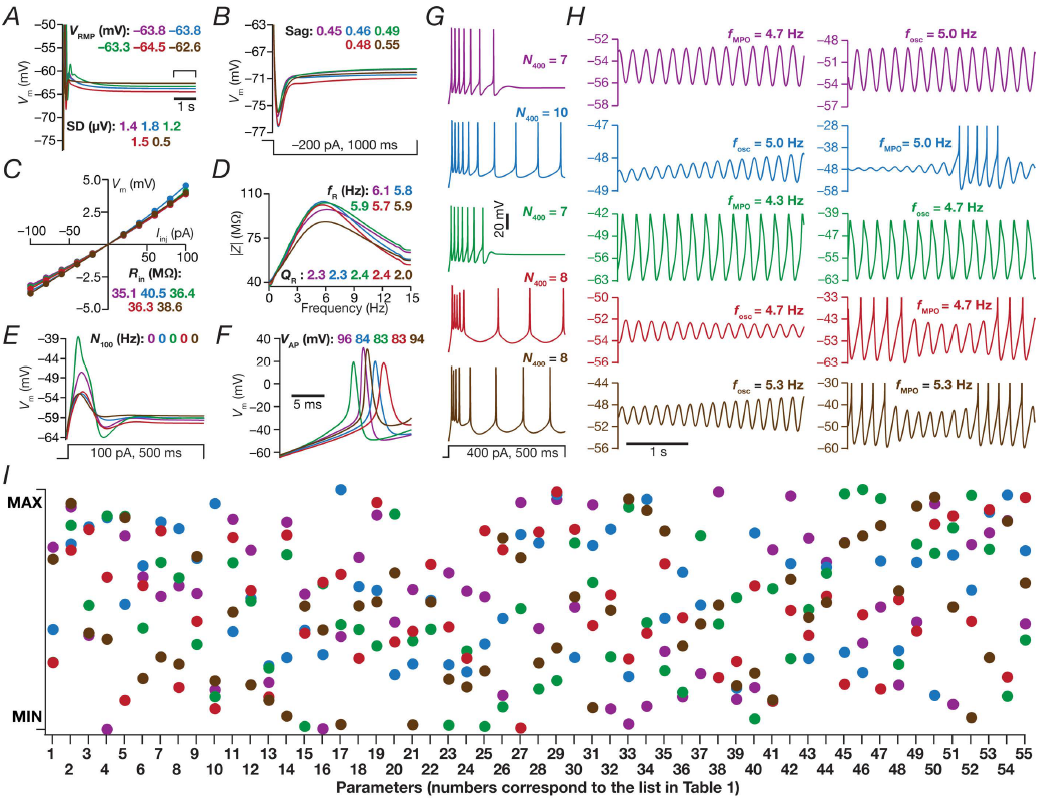
Disparate combinations of model parameters resulted in similar physiological measurements in 5 randomly chosen valid stellate cell models. (*A–H*) Voltage traces and 10 physiologically relevant measurements for 5 randomly chosen valid models obtained after MPMOSS. (*A*) Resting membrane potential (*V*_RMP_) and its standard deviation (SD), (*B*) Sag ratio, (*C*) Input resistance (*R*_in_), (*D*) Resonance frequency (*f*_R_) and resonance strength (*Q*_R_), (*E*) Number of action potentials for a step current injection of 100 pA for 500 ms (*N*_100_), (*F*) Amplitude of action potential (*V*_AP_), (*G*) Number of action potentials for a step current injection of 400 pA for 500 ms (*N*_400_) and (*H*) Peri-threshold membrane potential oscillation frequency (*f*_osc_). (*I*) Normalized values of each of the 55 parameters that were employed in the generation of stellate cell models, shown for the 5 randomly chosen models depicted in *A–H*. Each parameter was normalized by the respective minimum and maximum values that bound the stochastic search for that parameter (Table 1). Individual models are color coded across all plots and traces in *A–I*.

**Figure 5:**
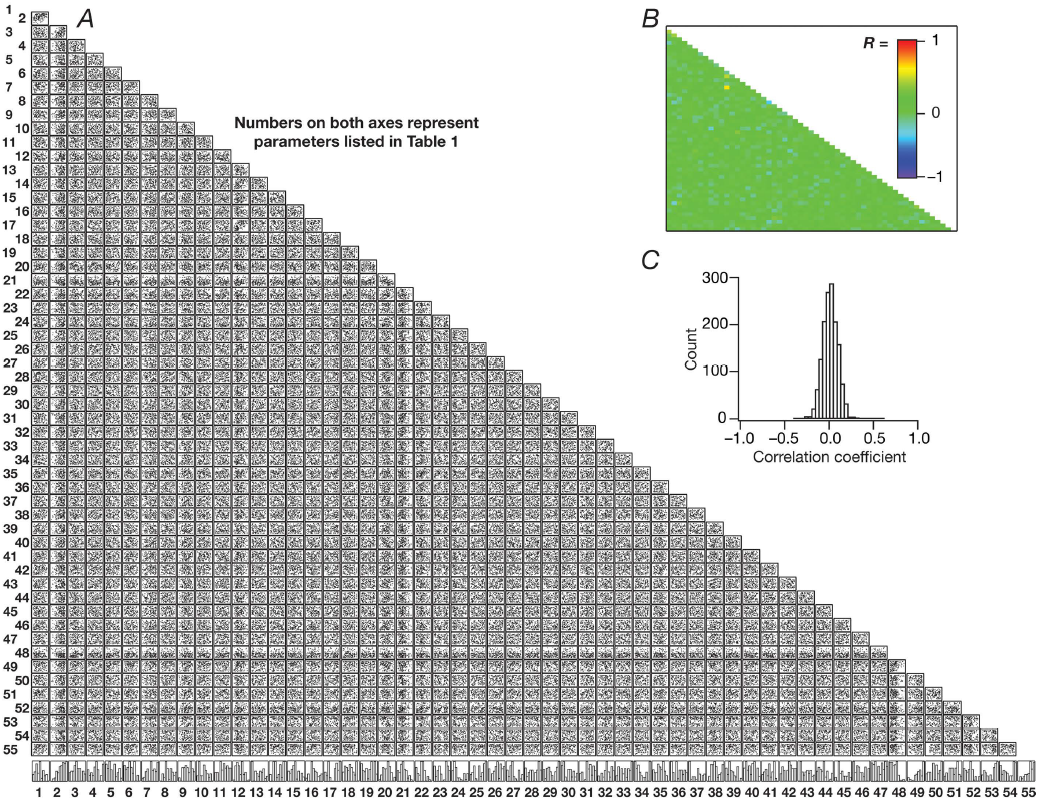
Expression of cellular scale degeneracy in the heterogeneous population of valid stellate cell models with weak pair-wise correlation among parameters. (*A*) Matrix consisting the pair-wise scatter plots between the 55 parameters (Table 1) for all 155 valid stellate cell models. Histograms in the bottom row depict the span of the corresponding parameter with reference to their respective min-max ranges (Table 1). (*B*) Heat map of the pair-wise correlation coefficient values corresponding to the scatter plots depicted in *A. (C)* Distribution of the 1485 unique correlation coefficient values from *B*.

Although the distributions of individual parameters span their respective ranges, was it essential that there are strong pair-wise constraints on different parameters towards obtaining valid models? To address this question and to test if parametric combinations resulting in the heterogeneous valid population were independent of each other, we plotted the pair-wise scatter plots across all 55 parameters for all the 155 valid models (Fig. 5A). We computed the Pearson's correlation coefficient for each pair-wise scatter plot (Fig. 5B), and found very weak correlations across different parameters (Figure 5B– C). Together, these results demonstrated that the ability of a heterogeneous model population to concomitantly match several electrophysiological signatures of LII SCs was attainable even when individual ion channels were expressed with disparate densities and properties and when channels did not express strong pairwise correlations. These provided strong lines of evidence for the expression of cellular-scale degeneracy in LII SCs, whereby disparate combinations of channels with distinct parameters were robustly capable of eliciting similar functional characteristics.

### Virtual knockout models: A many-to-many mapping between individual channels and physiological characteristics enabled the expression of degeneracy

A crucial requirement for the expression of such cellular-scale degeneracy is the ability of several channels to regulate different physiological characteristics (Drion *et al.*, 2015; Rathour *et al.*, 2016). In the absence of such capabilities, the system in essence will comprise of several one-to-one mappings between channels and physiological characteristics, thereby requiring the maintenance of individual channels at specific expression levels. In asking if there was a many-to-many mapping between channels and physiological properties, we employed virtual knockouts of individual channels on all models within the heterogeneous valid model population to assess the impact of their acute removal on physiology. We employed these virtual knockout models (VKMs) to assess the impact of individual channels on all the 10 physiological measurements by calculating the change observed in each measurement after setting individual conductance values to zero (Rathour & Narayanan, 2014; Anirudhan & Narayanan, 2015; Mukunda & Narayanan, 2017).

As expected, *V*_RMP_ (Fig. 6A) was largely unaffected by the knockout of supra-threshold conductances (KDR, NaF and HVA) with all subthreshold channels showing differential and heterogeneous regulation of *V*_RMP_. Specifically, consistent with prior electrophysiological recordings (Dickson *et al.*, 2000), HCN VKMs showed large and variable hyperpolarizing shifts in *V*_RMP_ with reference to their respective base valid models. In addition, knockout of KM or SK channels resulted in variable depolarizing shifts to *V*_RMP_, but VKMs of NaP, KA and LVA channels did not significantly alter *V*_RMP_. Although sag ratio was expectedly (Dickson *et al.*, 2000) reliant on HCN channels (Fig. 6B), there were other channels, including NaP, LVA, SK and KM channels, which also significantly contributed to the specific value of sag ratio. Input resistance (Fig. 6C) was critically altered by HCN channel knockouts (Dickson *et al.*, 2000), with other subthreshold channels including KM and SK also showing large impacts on *R*_in_.

**Figure 6:**
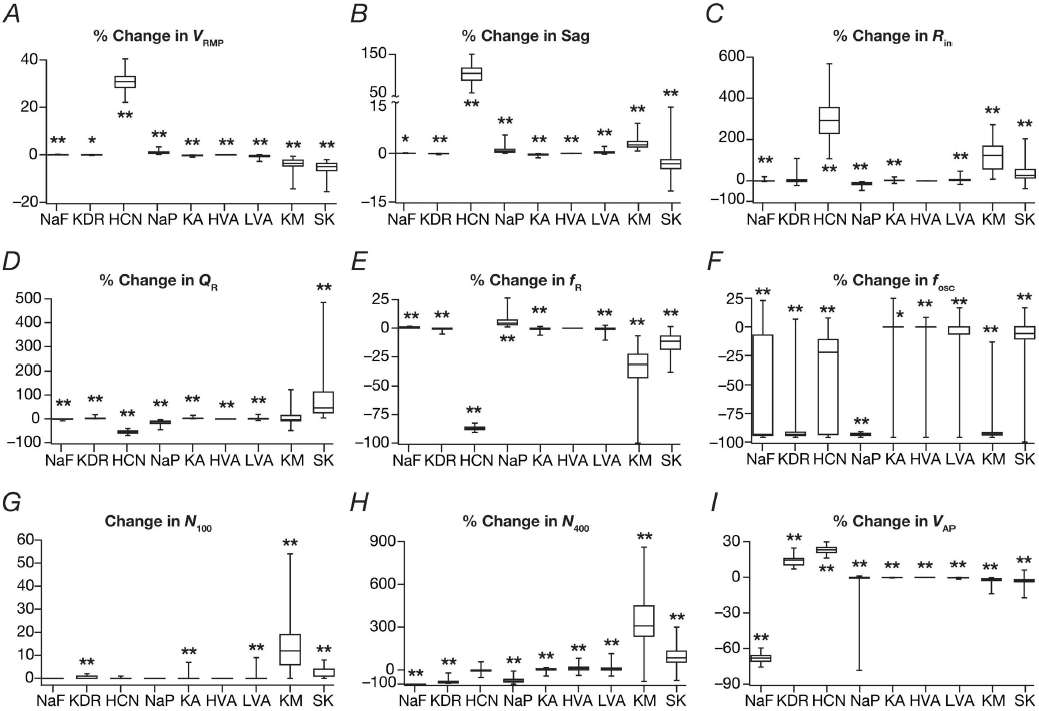
Single channel virtual knockout models (VKMs) unveiled differential and variable dependence of measurements on individual channels. (*A-I*) Change in different measurement values after virtual knockout of each channel from valid models obtained from the MPMOSS algorithm. Shown are percentage changes in resting membrane potential, *V*_RMP_ (*A*), sag ratio (*B*), input resistance *R*_in_ (*C*), resonance strength *Q*_R_ (*D*), resonance frequency *f*_R_ (*E*), peri-threshold oscillation frequency *f*_osc_ (*F*). Change in number of action potential elicited for 100 pA current injection (*N*_100_) is represented as a count (*G*) as *N*_100_ for all valid models was constrained to be zero, whereas changes in the number of action potentials elicited for 400 pA current injection *N*_400_ (*H*) and in action potential amplitude *V*_AP_ (*I*) are depicted as percentages. Note that VKMs that spontaneously fired or entered depolarization-induced block were removed from analyses owing to the inability to obtain subthreshold measurements. Consequently, for KDR knockouts *N*_VKM_ = 37, for KM knockouts *N*_VKM_ = 116, for SK knockouts *N*_VKM_ = 145 and all the other channel knockouts *N*_VKM_=155. For panels *A–I*, *: *p* < 0.01, **: *p* < 0.001, Wilcoxon rank sum test.

Resonance strength (Fig. 6D) and resonance frequency (Fig. 6E) were dramatically reduced in HCN knockouts (Erchova *et al.*, 2004; Boehlen *et al.*, 2013), with NaP, KM and SK channel knockouts also showing significant changes in both measurements. Of all the assessed measurements, we found the frequency of peri-threshold membrane potential oscillations to be the most sensitive measurement to channel knockouts, with most channels having a significant, yet variable, effect on *f*_osc_ (Fig. 6F). We observed the most reliable and least variable effect on *f*_osc_ with the removal of NaP, which consistently resulted in the loss of peri-threshold oscillations across all models (Fig. 6F). This is consistent with several studies that have demonstrated the importance of persistent sodium channels in the emergence of peri-threshold MPOs in LII SCs (Alonso & Llinas, 1989; Klink & Alonso, 1993; Dickson *et al.*, 2000; Boehlen *et al.*, 2013). Although NaP was the dominant channel that acted as the amplifying conductance (Hutcheon & Yarom, 2000) in the emergence of peri-threshold MPOs, we found heterogeneity across models in the specific resonating conductance that enabled these MPOs. Specifically, whereas in some models, the removal of HCN channels resulted in complete loss of MPOs, in other models the same result was achieved by the removal of KM channels. These observations suggested that the two conductances, HCN and KM, synergistically contributed as resonating conductances towards the emergence of peri-threshold MPOs in stellate cells (Nolan *et al.*, 2007; Boehlen *et al.*, 2013). Although most of the other channels showed regulatory capabilities in terms of regulating peri-threshold *f*_osc_, unlike NaP, HCN and KM channels, their removal did not result in complete elimination of MPOs in most models. The critical dependence of *f*_osc_ on KDR removal was just a reflection of high excitability of KDR VKMs, where the cells either spontaneously fired or abruptly switched to regular spiking with small current injections resulting in the complete absence of peri-threshold oscillations.

*N*_100_ was significantly higher with the deletion of KM or SK channels, with little or no effect with deletion of other channels (Fig. 6G). As spiking activity is critically reliant on KDR and NaF channels, their removal significantly reduced *N*_400_ (Fig. 6H). In addition, whereas the removal of the subthreshold regenerative conductance NaP reduced *N*_400_, knocking out the subthreshold restorative conductances, KM, SK and KA resulted in enhanced action potential firing that increased *N*_400_ to variable degrees (Fig. 6H). Although the two calcium channels mediate inward currents, their removal resulted in an increase (rather than a decrease) in *N*_400_ because of the presence of SK channels. Specifically, when either the HVA or the LVA channels was removed, the inward calcium current and cytosolic calcium concentration were lower, thereby resulting in lesser activation of SK channels and consequently leading to higher excitability (Fig. 6G– H). Finally, AP amplitude (Fig. 6I) was expectedly reliant on the presence of NaF channels, while KDR and HCN also had a regulatory role in setting the value of *V*_AP_. It should be noted that *V*_AP_ is significantly dependent on *V*_RMP_, as *V*_RMP_ determines the fraction of sodium channels that are inactivated and are thereby unavailable for activation. As the fraction of available sodium channels is higher with a hyperpolarized *V*_RMP_, *V*_AP_ is higher when *V*_RMP_ is hyperpolarized. As *V*_RMP_ was significantly hyperpolarized when HCN channels were knocked out (Fig. 6A), this implied that *V*_AP_ in HCN channel knockouts should be expected to be higher (Fig. 6I).

Together, analyses of all physiological measurements across valid models using VKMs of each of the 9 active ion channels demonstrated the clear lack of one-to-one mappings between channels and physiological characteristics. Although there was dominance of certain channels in their ability to alter specific measurements, there were several channels that were capable of regulating each measurement and each channel regulated several measurements. Additionally, the effect of virtually knocking out individual channels on all measurements was differential for different channels and measurements, and was variable for even a given channel-measurement combination. The electrophysiological support from LII SCs for several of our conclusions, including the regulatory role of specific channels and the differential and variable dependencies of measurements on channels, with reference to blockade of individual channels is strong (Alonso & Llinas, 1989; Klink & Alonso, 1993; Dickson *et al.*, 2000; Erchova *et al.*, 2004; Nolan *et al.*, 2007; Garden *et al.*, 2008; Pastoll *et al.*, 2012; Boehlen *et al.*, 2013). These observations together point to a many-to-many configuration of the mapping between channel properties and cellular-scale physiological characteristics, a critical substrate for neurons to exhibit cellular scale degeneracy (Figs. 3–5).

### Quantitative predictions: Spike initiation dynamics of stellate cells manifest theta-frequency spectral selectivity and gamma-band coincidence detection capabilities

As we now had a population of models that matched with LII SC physiology, we employed these models to make specific quantitative predictions about critical physiological characteristics of LII SC. Although it is well established that LII stellate cells exhibit spectral selectivity for subthreshold inputs (Haas & White, 2002; Erchova *et al.*, 2004; Giocomo *et al.*, 2007; Garden *et al.*, 2008; Giocomo & Hasselmo, 2008, 2009), it is not known if such frequency selectivity translates to the spike initiation dynamics as well (Das & Narayanan, 2014, 2015, 2017; Das *et al.*, 2017). To assess this, we employed zero-mean Gaussian white noise (GWN) with standard deviation adjusted such that the over all firing rate is ˜1 Hz (Fig. 7A). We computed the spike-triggered average (STA) as the mean of all the current stimuli (part of the GWN current, over a 300 ms period preceding each spike) that elicited a spike response in the model under consideration. We derived five distinct measurements of excitability, spectral selectivity and coincidence detection from the STA (Das & Narayanan, 2014, 2015, 2017; Das *et al.*, 2017), and repeated this for all the 155 valid models obtained from MPMOSS (Fig. 3–5).

**Figure 7:**
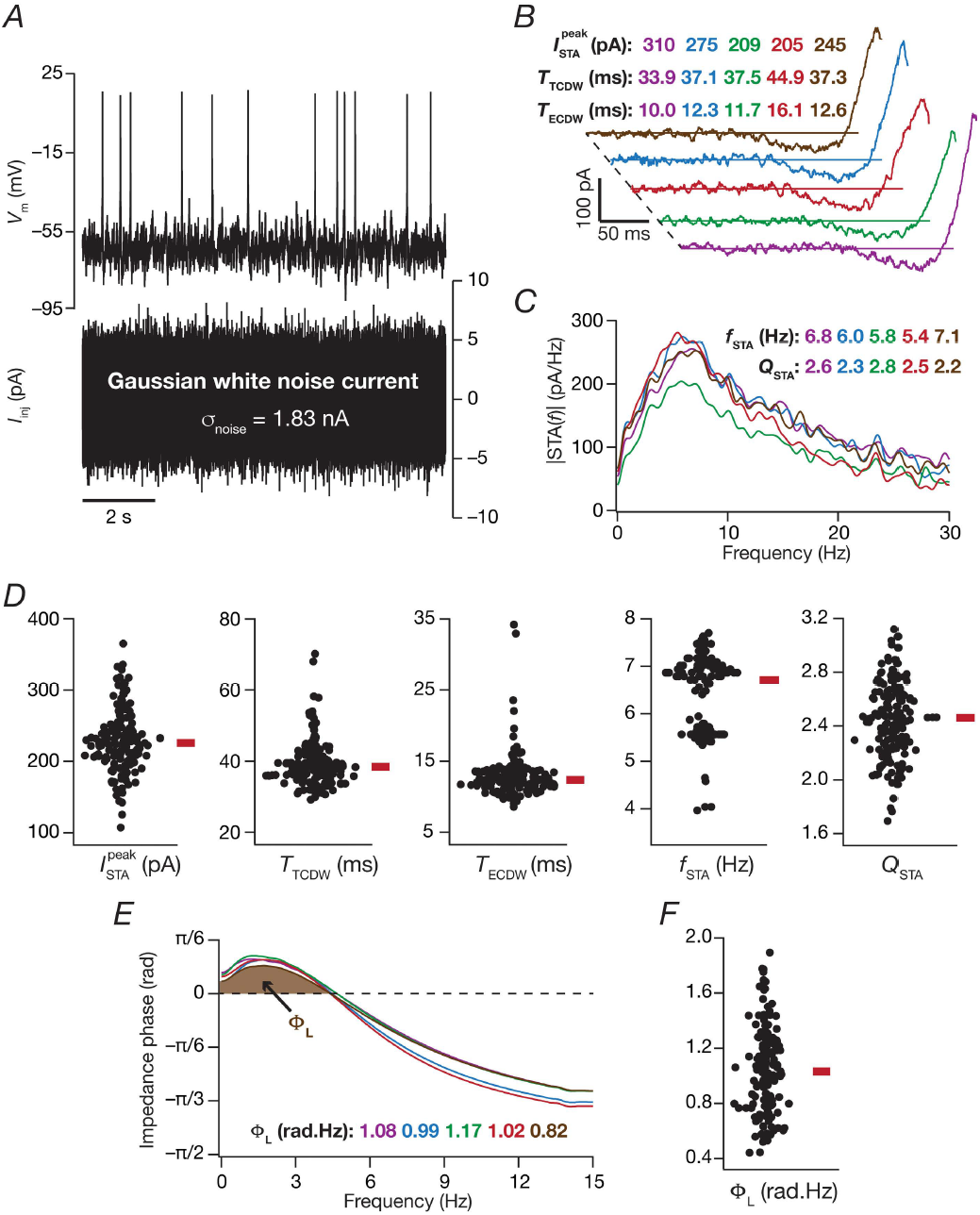
Measurements from the valid model population predict theta-frequency selectivity and gamma-range coincidence detection window in the spike triggered average of LII stellate cells. (*A*) *V*oltage response of an example valid model (top) to a zero-mean Gaussian white noise (GWN) current (bottom) of 10 s duration. σ_noise_= 1.83 nA. (*B–C*) Spike triggered average (STA) of the 5 valid stellate cell models shown in 5 randomly chosen valid stellate cell models shown in Fig. 4. Measurements derived from the temporal domain representation of STA were the peak STA current 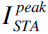, the total coincidence detection window (CDW) *T*_TCDW_ and the effective CDW *T*_ECDW_. (*C*) The magnitude of the Fourier transform of STA shown in *B*. Measurements derived from the spectral domain representation of STA were for the STA characteristic frequency *f*_STA_, and the strength of selectivity *Q*_STA_. *(D)* Bee swarm plots representing the distribution of the 5 quantitative measures of the STA for all 155 valid models. *(E)* Impedance phase profiles, and with values of total inductive phase, Φ_L_, defined as the area under the curve for the leading impedance phase (shaded portion) for 5 selected models. Color codes across this figure are matched with models in Fig. 4. *(F)* Distribution of Φ_L_ for all 155 valid stellate cell models.

The STA computed from the 5 models shown in Fig. 4 showed characteristic class II/III excitability (Fig. 7B), marked by the distinct presence of negative lobes in these STA (Haas & White, 2002; Ermentrout *et al.*, 2007; Haas *et al.*, 2007; Ratte *et al.*, 2013; Das & Narayanan, 2014, 2015, 2017; Das *et al.*, 2017). Employing quantitative metrics from the STA (Fig. 7B–C) for all the 155 valid models, we confirmed that these neurons were endowed with class II/III characteristics. Specifically, our analyses with the valid model population of LII SCs predict that the STA of these cells show theta-frequency spectral selectivity (Fig. 7D, *f*_STA_) with strong selectivity strength (Fig. 7D, *Q*_STA_). As class II/III excitability translates to coincidence detection capabilities in these neurons (Ratte *et al.*, 2013; Das & Narayanan, 2014, 2015, 2017; Das *et al.*, 2017), we computed two distinct measures of coincidence detection window (CDW) from the STA. Whereas the total CDW (*T*_TCDW_) considers the temporal span of the spike-proximal positive lobe of the STA, the effective CDW (*T*_ECDW_) also accounts for the specific shape of the STA in arriving at the CDW (Das & Narayanan, 2014, 2015, 2017). We computed these CDW measures for all the 155 models, and quantitatively predict that the LII SCs are endowed with gamma-range (25–150 Hz translating to 6.6–40 ms) coincidence detection capabilities (Fig. 7D). Finally, although it is known that the impedance phase of LII SCs manifest a low-frequency inductive lead (Erchova *et al.*, 2004), this inductive phase lead has not been systematically quantified. To do this, we computed the total inductive phase metric (Φ_L_; Fig. 7E) developed in (Narayanan & Johnston, 2008) for each of the 155 valid models, and quantitatively predict a prominent inductive phase lead in LII SCs (Fig. 8F) with specific values for Φ_L_.

**Figure 8:**
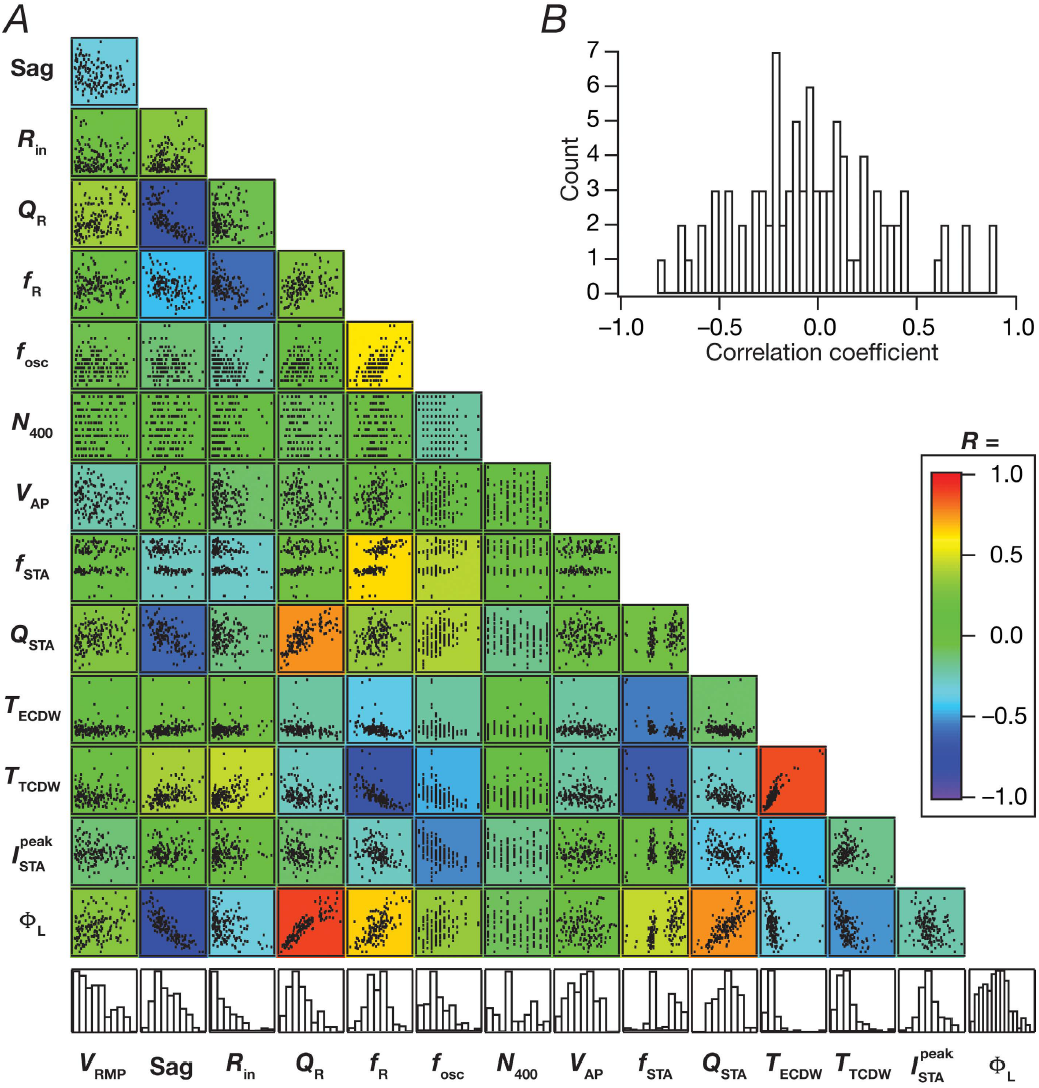
Pairwise correlations across physiological measurements from all valid stellate cell models were variable. *(A)* Matrix depicts the pair-wise scatter plots between the 14 measurements (8 physiologically relevant measurements, namely *V*_RMP_, Sag ratio, *R*_in_, *Q*_R_, *f*_R_, *f*_osc_, *N*_400_, *V*_AP_, in Fig. 1 and the 6 predicted measurements, namely 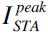, *T*_ECDW_, *T*_TCDW_, *f*_STA_, *Q*_STA_, Φ_L_, in Fig. 7). Histograms in the bottom row depict the span of the corresponding measurement with reference to their respective min-max ranges. Individual scatter plots are overlaid on a heat map that depicts the pair-wise correlation coefficient computed for that scatter plot, with the color code for the correlation coefficient values as provided. (B) Distribution of the 91 unique correlation coefficient values from scatter plots in *A*.

### Pair-wise correlations among measurements

Finally, as our analyses (Fig. 6) demonstrated a many-to-many mapping between biophysical parameters and physiological measurements, we asked if there were significant correlations among the distinct measurements. Strong correlations across these measurements (which are reflective of distinct physiological characteristics) would imply that they could be mapped to a smaller set of “core” measurements with the other measurements relegated to redundant and correlated reflections of these core measurements. In addition, strong correlations across measurements would also imply that there are measurements that are strongly reliant on the expression and properties of one specific channel. To assess correlations across measurements, we plotted the pairwise scatter plots (Fig. 8A) spanning all 14 measurements (8 measurements from Fig. 1: *V*_RMP_, Sag, *R*_in_, *Q*_R_, *f*_R_, *f*_osc_, *N*_400_, *V*_AP_ and 6 predicted measurements from Fig. 7: 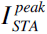, *T*_ECDW_, *T*_TCDW_, *f*_STA_, *Q*_STA_, Φ_L_) computed on the 155 valid models. We computed the Pearson's correlation coefficient for each scatter plot and analyzed the correlation coefficients across measurements (Fig. 8A–B). Although there were strong correlations across some measurements, a majority of these pair-wise correlations were weak (Fig. 8B). Measurements that showed strong pair-wise correlations were those that were known to be critically reliant on specific ion channels. For instance, sag ratio showed strong negative correlation with Φ_L_, *Q*_STA_ and *Q*_R_, whereas *f*_R_ manifested strong negative correlation with *R*_in_ and *T*_TCDW_, where all these measurements are known to have a strong dependence on HCN channels (Fig. 6). However, even within this subset of measurements that were strongly dependent on one channel, we noted that only a subset of the pair-wise correlations were high. For instance, with reference to the specific examples discussed above, *f*_R_ and *Q*_R_, both critically dependent on HCN channels and both derived from the impedance amplitude profile, do not show strong pair-wise correlation (Fig. 8A). Thus, although there were a small percentage of measurements that showed strong pair-wise correlations, a majority of their pair-wise correlations were weak, further emphasizing the absence of one-to-one relationships between channels and measurements (Fig. 6).

## DISCUSSION

The prime conclusion of this study is that LII stellate cells of the medial entorhinal cortex express cellular-scale degeneracy in the concomitant expression of several unique physiological characteristics of these neurons. We arrived at this conclusion through an unbiased stochastic search algorithm that spanned 55 parameters associated with biophysically constrained models of active and passive stellate cell components. We validated the outcomes of these stochastically generated models against 10 different physiological characteristics of stellate cells. This validation process provided us with a heterogeneous population of stellate cells that exhibited cellular-scale degeneracy with weak pair-wise correlations across parameters that governed these models. We employed these models to demonstrate the differential and variable dependencies of measurements on underlying channels, and also showed that the mapping between channels and measurements was many-to-many. Finally, we employed this electrophysiologically validated model population to make specific quantitative predictions that point to theta-frequency spectral selectivity and gamma-range coincidence detection capabilities in class II/III spike triggered average of LII SCs.

### Correlations between electrophysiological signatures of distal dendrites in CA1 pyramidal neurons and LII MEC stellate cells: Instances of cellular-scale efficient encoding?

A cursory glance at the electrophysiological properties of distal dendrites of CA1 pyramidal neurons and LII MEC stellate cells presents significant correlations between some physiological characteristics of these two structures. Although superficial layers of the MEC project to the distal dendrites of CA1 pyramidal neurons, it is LIII, and not LII, principal neurons of the MEC that project to CA1 pyramidal neurons (Andersen *et al.*, 2006). Despite this, there are several electrophysiological characteristics of LII MEC stellates that match with the distal dendrites of CA1 pyramidal neurons. Several of these similarities are strongly related to heavy expression profile of HCN channels in both these structures. Whereas the gradient in HCN channel density in CA1 pyramidal neurons implies a significantly sharp increase in HCN channels in the distal dendrites of CA1 pyramidal neurons (Magee, 1998; Lorincz *et al.*, 2002), the heavy expression of HCN channels in the cell body of LII MEC stellates is also well established (Klink & Alonso, 1993; Dickson *et al.*, 2000; Erchova *et al.*, 2004; Giocomo *et al.*, 2007; Nolan *et al.*, 2007; Garden *et al.*, 2008; Giocomo & Hasselmo, 2008, 2009; Giocomo *et al.*, 2011a; Boehlen *et al.*, 2013).

As a consequence of this, these two structures are endowed with significant sag, similar input resistances and comparable theta-band suthreshold resonance frequencies (Erchova *et al.*, 2004; Pastoll *et al.*, 2012). In addition to these, our study predicts that the STA of LII SCs should be endowed with theta-frequency spectral selectivity and gamma-band coincidence detection windows. Although it is known that the STA of LII SCs manifest class II/III excitability with coincidence detection characteristics (Haas & White, 2002; Haas *et al.*, 2007), quantification of spectral selectivity in the STA or a systematic assessment of the specific frequency band of coincidence detection capabilities has not been assessed. Our study specifically predicts the coincidence detection window (Fig. 7D; *T*_ECDW_) associated with the STA of LII SCs to be in the fast-gamma frequency (60–120 Hz) range, with a high theta-range spectral selectivity in the STA (Fig. 7D; *f*_STA_). Interestingly, quantitative predictions for these measurements for the distal dendritic region of CA1 pyramidal neurons also fall within the same spectral bands (Das & Narayanan, 2015).

This confluence of STA measurements of distal dendrites in CA1 pyramidal neurons and of LII SC soma, especially that of the coincidence detection window falling within the fast gamma-frequency band, is striking from the perspective of gamma-band multiplexing that has been reported in the CA1 subregion (Colgin *et al.*, 2009; Colgin & Moser, 2010; Fernandez-Ruiz *et al.*, 2017). Specifically, gamma oscillations in the CA1 subregion have been shown to manifest stratification into distinct fast and slow frequency components, impinging respectively on distal and proximal dendritic regions. This spatially stratified frequency-division multiplexing allows for differential coupling of CA1 pyramidal neurons to afferent inputs from the MEC and CA3 through different gamma bands. Juxtaposed against this, and within the efficient coding framework (Barlow, 1961; Bell & Sejnowski, 1997; Simoncelli & Olshausen, 2001; Lewicki, 2002; Simoncelli, 2003) where the response filters in a single neuron should match the natural statistics of afferent network activity (Narayanan & Johnston, 2012), it may be argued that different dendritic locations should be equipped with filters (STA kernels) that are matched to the afferent inputs (different gamma frequencies) that are received by that specific location. As a specific instance of such location-dependent efficient encoding of afferent network statistics in hippocampal pyramidal neurons, it has been quantitatively postulated that the distal dendrites of CA1 pyramidal neurons are endowed with coincidence detection windows specific to fast-gamma frequencies, whereas those of the proximal dendrites matched with slow-gamma frequencies (Das & Narayanan, 2015).

In this context and given the several lines of evidence for the dominance of fast gamma oscillations in the superficial layers of MEC (Colgin *et al.*, 2009; Colgin, 2016; Trimper *et al.*, 2017), our predictions that the CDW for stellate neurons should be in fast gamma band point to a similar form of efficient encoding schema in the MEC. Specifically, if the fast gamma oscillations are statistically the most prevalent in the superficial layers of MEC, it is imperative that neurons there are equipped with the machinery that is capable of detecting and processing inputs in this frequency range. Additionally, from the efficient coding perspective, as neurons tune their response properties to efficiently represent the statistics of afferent inputs (Wiesel & Hubel, 1963b, a; Hirsch & Spinelli, 1970; Bell & Sejnowski, 1997; deCharms *et al.*, 1998; Kilgard & Merzenich, 1998; Stemmler & Koch, 1999; Sharma *et al.*, 2000; Kilgard *et al.*, 2001; Simoncelli & Olshausen, 2001; Lewicki, 2002; Simoncelli, 2003; de Villers-Sidani *et al.*, 2007), it is essential that the response properties of LII MEC neurons are tuned to the fast gamma frequency range. Thus, our prediction on a fast gamma-band CDW in the STA of LII EC cells suggests the possibility of efficient encoding spanning the hippocampal formation, whereby the neuronal properties in terms of their class of excitability and specific band of frequency where their coincidence detection windows lie match with the type of gamma-frequency band that is most prevalent in that subregion (or strata in case of the CA1). A direct test of this experimental prediction would be to measure the CDW of pyramidal neurons in the CA3, of different dendritic subregions of the CA1 and of LII MEC stellates and ask if these CDW match with the respective gamma-band inputs that are prevalent in these subregions.

Finally, encoding schema are state-dependent processes that are critically reliant on behavioral state and consequent changes in afferent activity, neuromodulatory tones and activity-dependent plasticity (Marder, 2012; Narayanan & Johnston, 2012; Bargmann & Marder, 2013; Ratte *et al.*, 2013; Srikanth & Narayanan, 2015; Das *et al.*, 2017; Deneve *et al.*, 2017; Gallistel, 2017). Therefore, it is important that postulates on efficient codes and relationships between neuronal activity and afferent statistics are assessed in a manner that accounts for adaptability of coding within the neuron and across the network. Such activity-dependent plasticity and neuromodulation of intrinsic properties, especially of signature characteristics such as the membrane potential oscillations, could be systematically assessed in entorhinal stellates. Specifically, as activity-dependent plasticity of several ion channels is well established across different brain regions (Magee & Johnston, 2005; Narayanan & Johnston, 2007; Johnston & Narayanan, 2008; Narayanan & Johnston, 2008; Sjostrom *et al.*, 2008; Narayanan *et al.*, 2010; Shah *et al.*, 2010; Narayanan & Johnston, 2012), it would be important to ask if membrane potential oscillations, spectral selectivity characteristics (*f*_R_, *Q*_R_, *f*_STA_, *Q*_STA_, Φ_L_) and coincidence detection windows are amenable to such activity-dependent plasticity that target different ion channels (Fig. 6).

### Implications of cellular-scale degeneracy in LII stellate cell physiology

Degeneracy, the ability of disparate structural components to elicit similar function, is a ubiquitous biological phenomenon with strong links to robust physiology and evolution (Edelman & Gally, 2001; Price & Friston, 2002; Leonardo, 2005; Whitacre & Bender, 2010; Whitacre, 2010). Several studies spanning different systems have now demonstrated the expression of degeneracy, at different scales of analysis in neural systems as well (Marder & Prinz, 2002; Prinz *et al.*, 2004; Marder & Goaillard, 2006; Marder, 2011; Marder & Taylor, 2011; O'Leary & Marder, 2014; O'Leary *et al.*, 2014; Vogelstein *et al.*, 2014; Drion *et al.*, 2015). Within the hippocampal formation, recent studies have demonstrated the expression of degeneracy in single-neuron electrophysiology (Rathour & Narayanan, 2012; Srikanth & Narayanan, 2015; Mishra & Narayanan, 2017), intraneuronal functional maps (Rathour & Narayanan, 2014; Rathour *et al.*, 2016), synaptic localization required for sharp-tuning of hippocampal place fields (Basak & Narayanan, 2017), short-(Mukunda & Narayanan, 2017) and long-term (Anirudhan & Narayanan, 2015) synaptic plasticity profiles and network-scale response decorrelation (Mishra & Narayanan, 2017). In this study, we have demonstrated the expression of cellular-scale degeneracy in LII SCs, which are endowed with unique electrophysiological signatures including the prominent theta-frequency subthreshold membrane potential oscillations.

The implications for the expression of such cellular-scale degeneracy are several. First, the many-to-many mapping between channels and physiological characteristics (Fig. 6) and the consequent degeneracy in concomitantly achieving all signature electrophysiological characteristics implies that there is no explicit necessity for maintaining individual channels at specific levels or for maintaining paired expression between channel combinations (Figs. 4–5). This provides several significant degrees of freedom to the protein localization and targeting machinery in achieving specific functions or in maintaining homeostasis in these functional characteristics. Second, this also implies that adaptability to external stimuli, in terms of achieving efficient codes of afferent stimuli or in encoding features of a novel stimulus structure, could be achieved through disparate combinations of plasticity in several constituent components. For instance, our results predict that the ability to achieve fast gamma-band coincidence detection capabilities could be achieved through distinct combinations of channel parameters (Figs. 4–5, Fig. 7). Experimental analyses of such degeneracy in achieving efficient matching of neuronal response characteristics with the statistics of oscillatory patterns, through the use of distinct pharmacological agents that target different channels, would demonstrate the ability of different channels to regulate such efficient encoding (Das *et al.*, 2017). Finally, degeneracy in the expression of excitability properties also implies that similar long-term plasticity profiles in these neurons could be achieved with disparate combinations of parameters. Specifically, several forms of neuronal plasticity are critically reliant on the amplitude and kinetics of cytosolic calcium entry, which in turn are dependent on neuronal excitability properties. As similar excitability profiles could be achieved with distinct combinations of constituent components, it stands to reason that similar plasticity profiles could be achieved through disparate parameter combinations (Narayanan & Johnston, 2010; Ashhad & Narayanan, 2013; Anirudhan & Narayanan, 2015).

### Future directions: Electrophysiological and computational

A critical future direction for the study presented here is the incorporation of biological heterogeneities into entorhinal network models that assess grid cell formation. Most models for grid cell formation are simple rate-based models that are made of homogeneous repeating units, and even models that incorporate conductance-based neurons for grid cell modeling do not account for the several biological heterogeneities that are expressed in the entorhinal network. This lacuna is especially striking because such analysis is critical for the elucidation of the mechanistic bases for grid cell formation (in terms of the channels and receptors involved) and for the quantitative understanding of the ability of the entorhinal network to elicit robust grid cell behavior in the presence of several network heterogeneities. For instance, are the several forms of network heterogeneities (*i.e.*, intrinsic, local synaptic and afferent connectivity) aiding or hampering the robustness of grid cell emergence? How do different forms of heterogeneities interact in the manifestation of the specific neuronal and network properties that are essential for grid cell formation? Are the different models for grid cell emergence robust to significant variability in channels, synapses and afferent connectivity? How do different channels and receptors, within the limits of biological variability in their expression and properties, contribute to the emergence of grid cells? Are specific forms of neuronal and network heterogeneities essential for grid cell formation? How are the different signature electrophysiological characteristics of the entorhinal neurons critical in the formation of grid cells? Does class II/III excitability of LII stellate cells play a critical role in entorhinal function and in grid cell formation?

At the cellular scale, future studies could build heterogeneous models to account for the signature continuum of intrinsic physiological characteristics along the dorsoventral axis of the MEC, also accounting for specific channels that are known to change along this axis (Giocomo *et al.*, 2007; Garden *et al.*, 2008; Giocomo & Hasselmo, 2008, 2009; Yoshida *et al.*, 2011). Such studies will provide quantitative bases for exploring the expression of degeneracy in maintaining the dorsoventral gradients, and could be incorporated into network models for grid formation in assessing the relationship between grid-cell characteristic and neuronal intrinsic properties.

Additionally, the heterogeneous model population built in this study comprises a simple single-compartmental structure that did not account for dendritic properties or morphological heterogeneity of LII SCs. Electrophysiologically, the absence of systematic cell-attached recordings of specific channels and their properties in the stellate cells has been a significant impediment in building morphologically realistic models. While future experimental studies could focus on recording channels and channel properties along the non-planar dendritic arbor of stellate cells, future computational studies could incorporate these channels into morphologically realistic models to assess the specific roles of dendritic channels and morphological heterogeneity in grid cell formation (Schmidt-Hieber *et al.*, 2017). Finally, our study has specific quantitative predictions about the STA of these neurons and also presents an array of cross-dependencies of measurements on different channel types. Future electrophysiological studies could systematically test these predictions, and assess efficient encoding in these structures apart from adding further evidence for the many-to-many mapping between channels and physiological characteristics. For instance, an important prediction from VKMs is on the critical role of SK channels in regulating several intrinsic measurements including membrane potential oscillations (Fig. 6). Although the expression of calcium-dependent potassium channels is established in stellate cells (Khawaja *et al.*, 2007; Pastoll *et al.*, 2012), the specific role of these channels in regulating resonance, impedance phase, intrinsic excitability and membrane potential oscillations could be tested electrophysiologically using pharmacological blockers of SK channels.

## AUTHOR CONTRIBUTIONS

D.M. and R. N. designed experiments; D.M. performed experiments and carried out data analysis; D.M. and R. N. co-wrote the paper.

## ACKNOWLEDGMENTS

The authors thank members of the cellular neurophysiology laboratory for helpful discussions and for comments on a draft of this manuscript.

